# A neuroendocrine pathway modulating osmotic stress in *Drosophila*

**DOI:** 10.1101/522441

**Authors:** Meet Zandawala, Thomas Nguyen, Marta Balanyà Segura, Helena A. D. Johard, Mirjam Amcoff, Christian Wegener, Jean-Paul Paluzzi, Dick R. Nässel

## Abstract

Environmental factors challenge the physiological homeostasis in animals, thereby evoking stress responses. Various mechanisms have evolved to counter stress at the organism level, including regulation by neuropeptides. In recent years, much progress has been made on the mechanisms and neuropeptides that regulate responses to metabolic/nutritional stress, as well as those involved in countering osmotic and ionic stresses. Here, we identified a peptidergic pathway that links these types of regulatory functions. We uncover the neuropeptide Corazonin (Crz), previously implicated in responses to metabolic stress, as a neuroendocrine factor that inhibits the release of a diuretic hormone, CAPA, and thereby modulates the tolerance to osmotic and ionic stress. Both knockdown of *Crz* and acute injections of Crz peptide impact desiccation tolerance and recovery from chill-coma. Mapping of the Crz receptor (*CrzR*) expression identified three pairs of *Capa-*expressing neurons (Va neurons) in the ventral nerve cord that mediate these effects of Crz. We show that Crz acts to restore water/ion homeostasis by inhibiting release of CAPA neuropeptides via inhibition of cAMP production in Va neurons. Knockdown of *CrzR* in Va neurons affects CAPA signaling, and consequently increases tolerance for desiccation, ionic stress and starvation, but delays chill-coma recovery. Optogenetic activation of Va neurons stimulates excretion and simultaneous activation of Crz and CAPA-expressing neurons reduces this response, supporting the inhibitory action of Crz. Thus, Crz inhibits Va neurons to maintain osmotic and ionic homeostasis, which in turn affects stress tolerance. Earlier work demonstrated that systemic Crz signaling restores nutrient levels by promoting food search and feeding. Here we additionally propose that Crz signaling also ensures osmotic homeostasis by inhibiting release of CAPA neuropeptides and suppressing diuresis. Thus, Crz ameliorates stress-associated physiology through systemic modulation of both peptidergic neurosecretory cells and the fat body in *Drosophila*.

**Author summary:** Insects are among the largest groups of animals and have adapted to inhabit almost all environments on Earth. Their success in surviving extreme conditions stems largely from their ability to withstand environmental stress, such as desiccation and cold. However, the neural mechanisms that are responsible for coordinating responses to counter these stresses are largely unknown. To address this, we delineate a neuroendocrine axis utilizing the neuropeptides Corazonin (Crz) and CAPA, that coordinate responses to metabolic and osmotic stress. We show that Crz inhibits the release of a diuretic peptide, CAPA from a set of neurosecretory cells. CAPA in turn influences osmotic and ionic balance via actions on the Malpighian tubules (the insect analogs of the kidney) and the intestine. Taken together with earlier work, our data suggest that Crz acts to restore metabolic homeostasis at starvation and osmotic homeostasis during desiccation by inhibiting release of the diuretic hormone CAPA. Hence, this work provides a mechanistic understanding of the neuroendocrine mitigation of metabolic and osmotic stress by two peptide systems.

## Introduction

Environmental conditions continuously challenge the physiological homeostasis in animals, thereby evoking stress that can adversely affect the health and lifespan of an individual. For instance, lack of food and water, extreme temperatures, infection and predation can all evoke stress responses. In order to counter this stress and restore homeostasis, animals have evolved a multitude of physiological and behavioral mechanisms, which involve actions of multiple tissues and/or organs (see 1, 2). The core of these mechanisms involves hormones and neuropeptides, which orchestrate the actions of various organs to counteract stress and maintain homeostasis. One well studied mechanism counteracting water-deficit stress is the mammalian anti-diuretic system that involves hypothalamic osmoreceptors stimulating the sensation of thirst that leads to the release of the anti-diuretic hormone vasopressin, which targets multiple organs, including the kidney, to decrease urine output and conserve water (3, 4). In insects such as the vinegar fly, *Drosophila melanogaster*, much progress has been made on the mechanisms and factors regulating metabolic homeostasis, nutritional stress and longevity (see 1, 5–8). Several neuropeptides and peptide hormones have been shown to influence responses to nutrient stress via actions on peripheral tissues such as the liver-like fat body. Specifically, these peptide hormones include *Drosophila* insulin-like peptides (DILPs), adipokinetic hormone (AKH) and corazonin (Crz) (9–16). In addition, mechanisms regulating the release of these hormones are being unraveled (12, 17, 18). Hence, we begin to understand the neural circuits and neuroendocrine pathways regulating metabolic homeostasis, and nutritional stress. However, the circuits and/or pathways that regulate thermal, osmotic and ionic stresses remain largely unexplored.

Thus, we ask what factors and cellular systems constitute the osmoregulatory axis in *Drosophila* (and other insects)? Since nutrient and osmotic homeostasis are inter-dependent (19, 20), we hypothesized that regulation of osmotic stress may involve factors that also regulate nutritional stress. Here, we identified Crz signaling, a paralog of the AKH/Gonadotropin-releasing hormone signaling system (21, 22), as a candidate regulating these stresses. Based on previous research in *Drosophila* and other insects, it has been hypothesized that Crz modulates responses to stress, especially nutritional stress (16, 23, 24). Consistent with this, recent work has shown that neurosecretory cells co-expressing Crz and short neuropeptide F (sNPF) are nutrient sensing (17, 25) and modulate nutrient homeostasis through differential actions of the two co-expressed neuropeptides. Whereas sNPF acts on the insulin-producing cells (IPCs) and AKH-producing cells to stimulate DILP release and inhibit AKH release, respectively, systemic Crz signaling modulates feeding and nutritional stress through actions on the fat body (9, 14, 17).

Although the role of Crz in regulating responses to nutritional stress is now established, less is known about its role in cold tolerance and ion/water homeostasis. Hence, to address this, we examine the role of Crz in modulating osmotic and ionic stresses and furthermore outline the cellular systems constituting the Crz signaling axis. To this end, we analyzed the effects of manipulating Crz signaling on desiccation tolerance and chill-coma recovery as these two assays are routinely used to assess responses to osmotic/ionic stress (26, 27). Both knockdown of *Crz* and acute injections of Crz peptide impact desiccation tolerance and recovery from chill-coma. Comprehensive mapping of the Crz receptor (*CrzR*) expression revealed that these effects of Crz are not likely mediated by direct modulation of the osmoregulatory tissues but indirectly via three pairs of neurons (Va neurons) in the ventral nerve cord (VNC), which express diuretic CAPA neuropeptides. Knockdown of the *CrzR* in Va neurons affects CAPA release and ion/water balance, consequently influencing desiccation tolerance and chill-coma recovery. Crz acts via inhibition of cAMP production in Va neurons to inhibit release of CAPA and thereby reducing water loss during desiccation. Our data, taken together with published findings (14) suggest that Crz is released into the hemolymph during nutritional stress and acts on the fat body to mobilize energy for food search to increase food intake. In summary, we propose that Crz acts upstream of CAPA signaling to regulate water and ion balance as well as restore nutrient levels caused by starvation. Thus, in addition to the hormonal actions of Crz on the fat body to maintain metabolic homeostasis and counter nutritional stress (14), this peptide also helps maintain osmotic homeostasis.

## Results

### *Crz* knockdown influences osmotic and ionic stresses

In adult *Drosophila*, Crz is expressed in two major cell clusters (28): dorsal lateral peptidergic neurons (DLPs) in the pars lateralis of the brain **(S1A Figure)** which co-express sNPF (9) and 2-3 pairs of male-specific interneurons in the abdominal neuromeres of the VNC **(S1B Figure)** (29). The DLPs can be further subdivided into two groups; a pair of large neurosecretory cells and several pairs of smaller neurons in the superior lateral protocerebrum, some of which are neurosecretory while others innervate the antennal lobe **(S1C Figure)**. Previously, we showed that global knockdown of *Crz* (*Crz>Crz-RNAi*) increased survival of flies exposed to desiccation (14). Hence, we asked whether global knockdown of *Crz* also influences chill-coma recovery which indicates dysregulation of the ionic and osmotic homeostasis of the fly. Indeed, knockdown of *Crz* using *Crz-RNAi*^*#1*^ **(Figure 1A)** also results in delayed recovery from chill-coma **(Figure 1B)** and also increased survival under ionic stress **(Figure 1C)**. Since an independent *Crz-RNAi*^*#2*^ was ineffective at knocking down *Crz* **(S2A-B Figures)**, we did not use it for subsequent experiments. In addition, adult-specific pan-neuronal knockdown of *Crz* using the GeneSwitch system (30) also resulted in increased survival under ionic stress **(S2C Figure)**. Taken together, these experiments indicate that Crz influences ionic and osmotic stresses.

**Figure 1:**
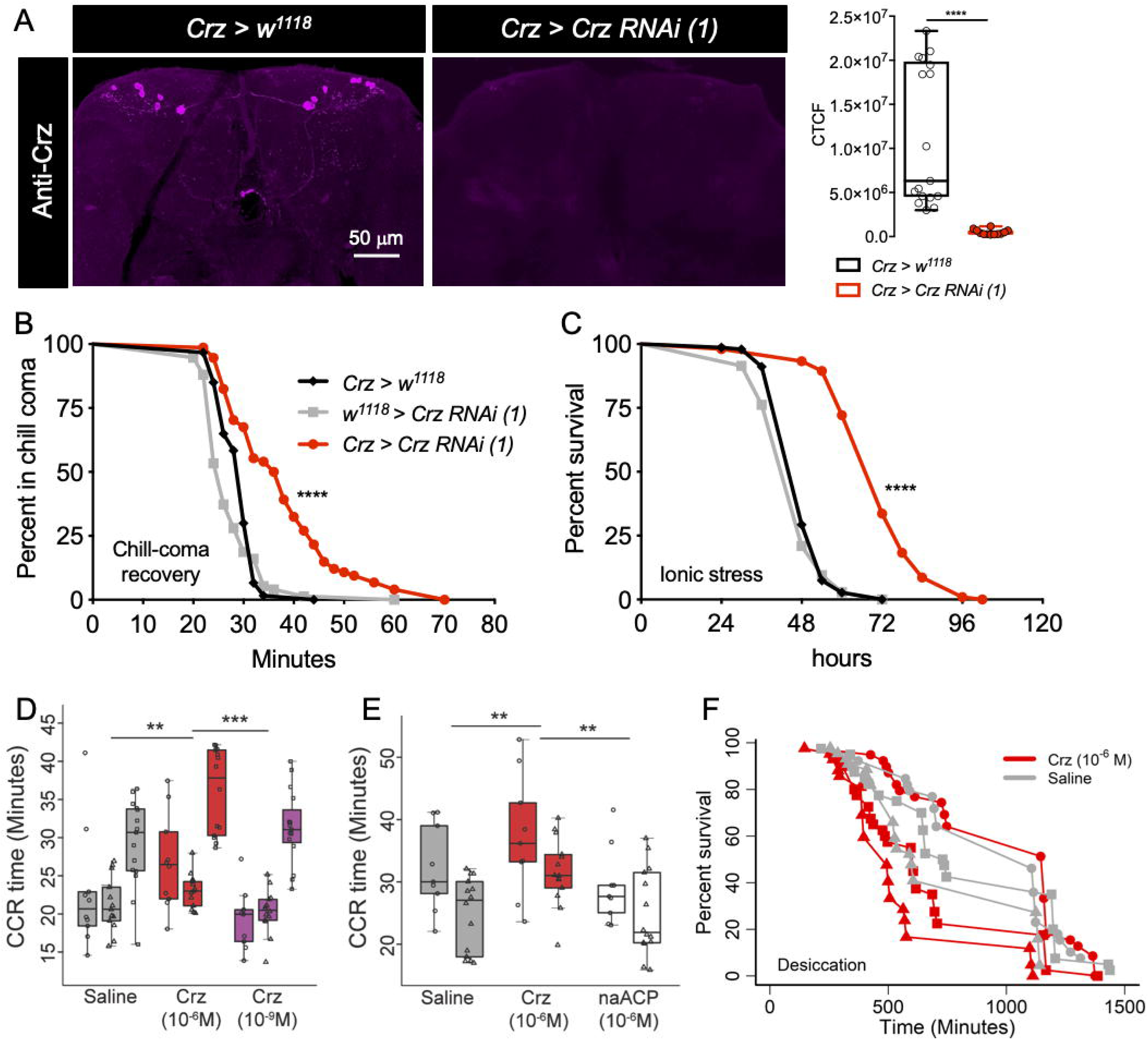
Crz signaling influences osmotic and ionic stresses. **(A)** *Crz^1^-GAL4* driven *Crz-RNAi (1)* causes a significant decrease in anti-Crz staining (corrected total cell fluorescence, CTCF) in the brains of adult *Drosophila* (**** p < 0.0001 as assessed by unpaired *t-*test). **(B)** *Crz* knockdown results in a significant delay in recovery from chill-coma and **(C)** increased survival under ionic stress. For **B** and **C**, data are presented as survival curves (**** p < 0.0001, as assessed by Log-rank (Mantel-Cox) test). Crz peptide injections *in vivo* influence **(D, E)** chill-coma recovery and **(F)** desiccation survival. **(D)** Low dose (10^−9^M) of injected Crz has no effect whereas a high dose (10^−6^M) of Crz delays chill-coma recovery (** p < 0.01, *** p < 0.001 as assessed by two-way ANOVA). **(E)** A non-active exogenous peptide (10^−6^M), non-amidated *Aedes* adipokinetic hormone/corazonin-related peptide (naACP), has no effect on chill-coma recovery (** p < 0.01 as assessed by two-way ANOVA). In **D** and **E**, the y-axes (Time) represents the chill-coma recovery (CCR) time. **(F)** We used a Cox proportional hazards model to test whether injection of 10^−6^M Crz (red lines) influences the rate of death under desiccation conditions compared to saline injection (grey lines), while accounting for any time-effects (different time points are indicated using circle, triangle and square symbols). Crz injection increases the risk of death by 40% compared to control flies. This difference is statistically significant at p=0.0003. Two independent trials for **E** and three independent trials for **D** and **F** were performed. These trials are indicated using different symbols (circle, triangle and square).

### Crz peptide injections influence desiccation survival and chill-coma recovery

Having shown that chronic genetic manipulations that interfere with Crz signaling impacts stress tolerance, we asked whether acute increase of Crz signaling also affects these stress responses. To this end, we first quantified the recovery time of flies from chill-coma that had been injected with Crz. An injection of a high dose of Crz (10^−6^M), but not a low dose (10^−9^M), delays recovery from chill-coma **(Figure 1D)**. Injection of an inactive exogenous peptide (non-amidated *Aedes aegypti* ACP, naACP) has no effect on chill coma recovery **(Figure 1E)**, suggesting that these effects are not due to changes in hemolymph osmolarity following peptide injection. Next, we independently assayed the Crz-injected flies for survival under desiccation conditions. Flies injected with 10^−6^M Crz display decreased desiccation tolerance compared to saline-injected flies **(Figure 1F)**. Taken together, these results suggest that Crz decreases resistance to desiccation and cold exposure.

### *CrzR* is expressed in CAPA and other peptidergic neurons

Since Crz signaling influences osmotic and ionic stresses, we wanted to outline the pathways mediating this response. We predicted that Crz may modulate these stresses via either directly acting on peripheral tissues such as the Malpighian tubules (analogous to the human kidney) and/or hindgut and rectum that regulate ionic and water homeostasis, or indirectly through one (or more) of the diuretic/anti-diuretic peptide hormones that act on these organs. To determine the mode of Crz action, we examined expression of the *CrzR* in the CNS and peripheral tissues by driving GFP expression using different *CrzR-GAL4* lines. *CrzR-GAL4*^*T11a*^ and *CrzR-GAL4*^*Se*^ had similar expression patterns; however, *CrzR-GAL4*^*T11a*^ resulted in stronger GFP expression than *CrzR-GAL4*^*Se*^. Hence, the *CrzR-GAL4*^*T11a*^ line (referred to as *CrzR-GAL4* from hereon) was used for all the subsequent experiments. In the periphery, *CrzR* is not expressed in the midgut, hindgut or Malpighian tubules **(S3A Figure)**. However, weak *CrzR* expression is present in sparse efferent neuronal projections to the rectal pad **(S3B Figure)**. Overall, this is consistent with the RNA-Seq expression of *CrzR* reported on FlyAtlas 2 (http://flyatlas.gla.ac.uk/) **(S3C Figure)**; however, these findings suggest that the effects of Crz on diuresis, and thus ionic and osmotic stress tolerance, are likely mediated via another neuropeptide and not by direct actions on peripheral osmo- and ionoregulatory tissues.

In *Drosophila*, there are four main neuropeptides (primarily diuretic) that are known to influence water and ionic homeostasis via direct actions on peripheral tissues: CAPA, diuretic hormone 44 (DH44), diuretic hormone 31 (DH31) and leucokinin (LK) (see 6). As a first step to determine if any of these neuropeptides act downstream of Crz signaling, we comprehensively mapped the distribution of *CrzR* in the adult CNS and co-labelled these specimens with antibodies against various neuropeptides. Within the adult brain, *CrzR* is expressed in distinct cell clusters **(S4A Figure)** and branches of these neurons partly superimpose the Crz-expressing DLP neuron processes **(S4B-C Figures)**. In the adult VNC, the *CrzR* is strongly expressed in three pairs of neurons in the abdominal neuromeres A2-A4 **(S4D Figure)**. In addition, weak GFP expression is also observed in two pairs of posterior neurons (indicated by a white arrow). These might be the efferent neurons that innervate the rectal pad **(S3B Figure)**. Using peptide immunolabeling we found that *CrzR* is expressed in a single pair of neurons in the SEZ and three pairs of Va neurons in the VNC, which express the *Capa* gene (*Capa* encodes CAPA-1, CAPA-2 and pyrokinin-1 (PK-1) neuropeptides) **(Figure 2A-B)** (31–33). In addition, *CrzR* is weakly expressed in a subset of the six DH44-expressing median neurosecretory cells in the pars intercerebralis **(S5 Figure)**. However, *CrzR-GAL4* driven GFP expression was not detected in neurons expressing DH31 **(S6A-C Figures)** or LK **(S6D-F Figures)**. CAPA peptides produced by the Va neurons have been previously implicated in affecting tolerance to desiccation and cold via actions on the Malpighian tubules (26). Moreover, as a result of differential prohormone processing, the *Capa* gene expressing neurons in the SEZ only produce a pyrokinin-1 neuropeptide, whereas the Va neurons can produce all three peptides encoded by *CAPA* (33). Since DH44 knockdown has no effect on desiccation tolerance (34), and since we detected strong *CrzR* expression in Va neurons but not DH44 neurons, we focus here on the Crz-CAPA pathway, specifically the CAPA-expressing Va neurons. To further validate *CrzR* expression in Va neurons, we utilized a recently generated *CrzR-T2A-GAL4* line where the T2A-GAL4 encoding cassette is inserted immediately upstream of the stop codon of *CrzR* (35). As expected, *CrzR-T2A-GAL4* also drove GFP expression in Va neurons (**Figure 2C**) further supporting the link between Crz and CAPA. Lastly, single-cell transcriptome datasets of the adult fly (36, 37) also reveal that *CrzR* and *Capa* are co-expressed in the brain and VNC **(Figure 2D)** further confirming *CrzR* expression in *Capa-* expressing neurons.

**Figure 2:**
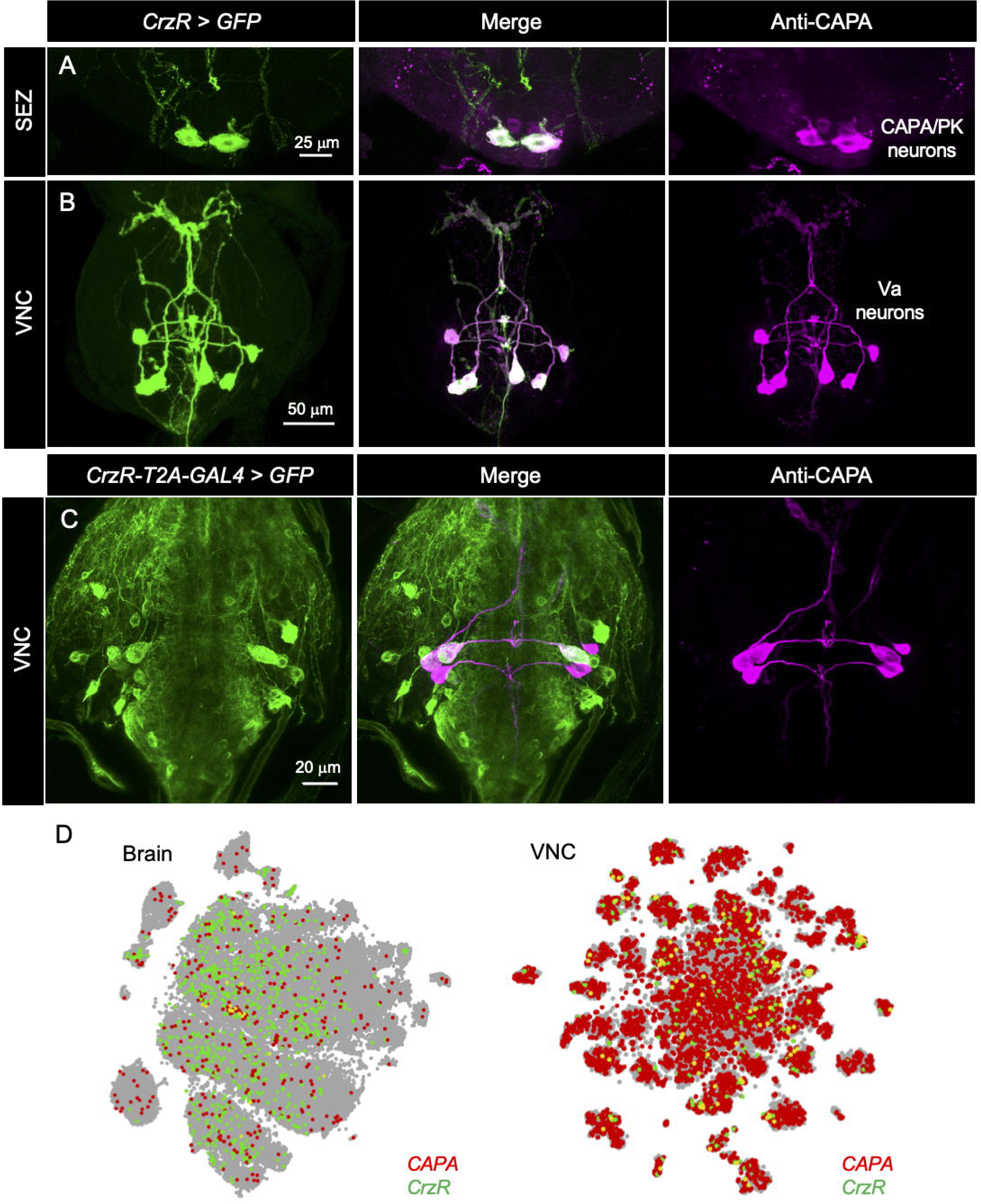
CrzR is expressed in CAPA neurons. *CrzR-GAL4* drives GFP expression in CAPA/pyrokinin (CAPA/PK) producing neurons (labeled with anti-CAPA antibody) in **(A)** the subesophageal zone (SEZ) and **(B)** ventral nerve cord (VNC) of adult males. The three pairs of neurons in the VNC are referred to as Va neurons. **(C)** *CrzR-T2A-GAL4* also drives GFP expression in Va neurons. **(D)** Mining the single-cell transcriptome data sets reveals that *Capa* and *CrzR* as coexpressed in the brain and VNC of adult *Drosophila*. Data was mined using Scope (http://scope.aertslab.org) (36, 37).

Additionally, we tried to identify potential interactions between Crz and other neurons producing feeding and stress-regulating peptides using a similar approach. The *CrzR* is not expressed in insulin-producing cells **(S7A Figure)** or AKH-producing cells **(S7B Figure)**, consistent with previous data identifying sNPF as the functional neurotransmitter in DLPs affecting DILP and AKH release (17). Interestingly, *CrzR* is expressed in a subset of the *Hugin*-producing neurons (labelled using anti-PRXa antibody) (38) in the SEZ **(S7C Figure)**, as well as neurons in the SEZ reacting with antiserum to FMRFamide (this antiserum cross reacts with neuropeptides such as sNPF, NPF, sulfakinin, FMRFamide and myosuppressin, which all have an RFamide C-terminus) **(S7D Figure)**. Furthermore, *CrzR* was detected in PDF-expressing small ventrolateral clock neurons (sLNvs) (39) **(S7E-F Figures)**. The Hugin neuropeptide (Hugin-PK) has been shown to regulate feeding and locomotion in larval and adult *Drosophila* (40–42) and PDF-expressing sLNvs are part of the circadian clock circuit (43, 44).

### Crz inhibits cAMP levels in Va neurons

Having shown that *CrzR* is expressed in Va neurons, we asked whether Va neurons can respond directly to Crz. Since the identity of the second messengers utilized by CrzR is unknown, we first predicted the coupling specificity of CrzR to various G-proteins (http://athina.biol.uoa.gr/bioinformatics/PRED-COUPLE2/) (45). This analysis revealed high coupling specificity of CrzR to Gi/o G-protein (p-value = 1.69 × 10^−6^) suggesting that Crz may inhibit the cAMP dependent pathway. Consistent with this prediction, bath application of Crz to dissected VNC diminished NKH477 (a forskolin analog and potent adenylyl cyclase activator) stimulated cAMP levels in Va neurons, indicated by the response of the FRET-based cAMP sensor Epac1-camps expressed using *CAPA-GAL4* **(Figures 3A-D)**. Addition of Crz alone had no impact on the cAMP FRET signal compared to saline controls **(Figures 3A-B)**. However, either an addition of NKH477 to preparations pre-incubated with Crz **(Figures 3A-B)** or simultaneous coapplication of Crz and forskolin **(Figures 3C-D)** resulted in a reduced cAMP FRET signal compared to their respective controls. In addition, Crz application to *ex vivo* VNC preparations of *CAPA > GCaMP6m* flies had no impact on Ca^2+^ levels in Va neuron as measured using GCaMP fluorescence **(Figures 3E-F)**. These results suggest that Crz inhibits cAMP levels in Va neurons, most likely by direct action on the CrzR.

**Figure 3:**
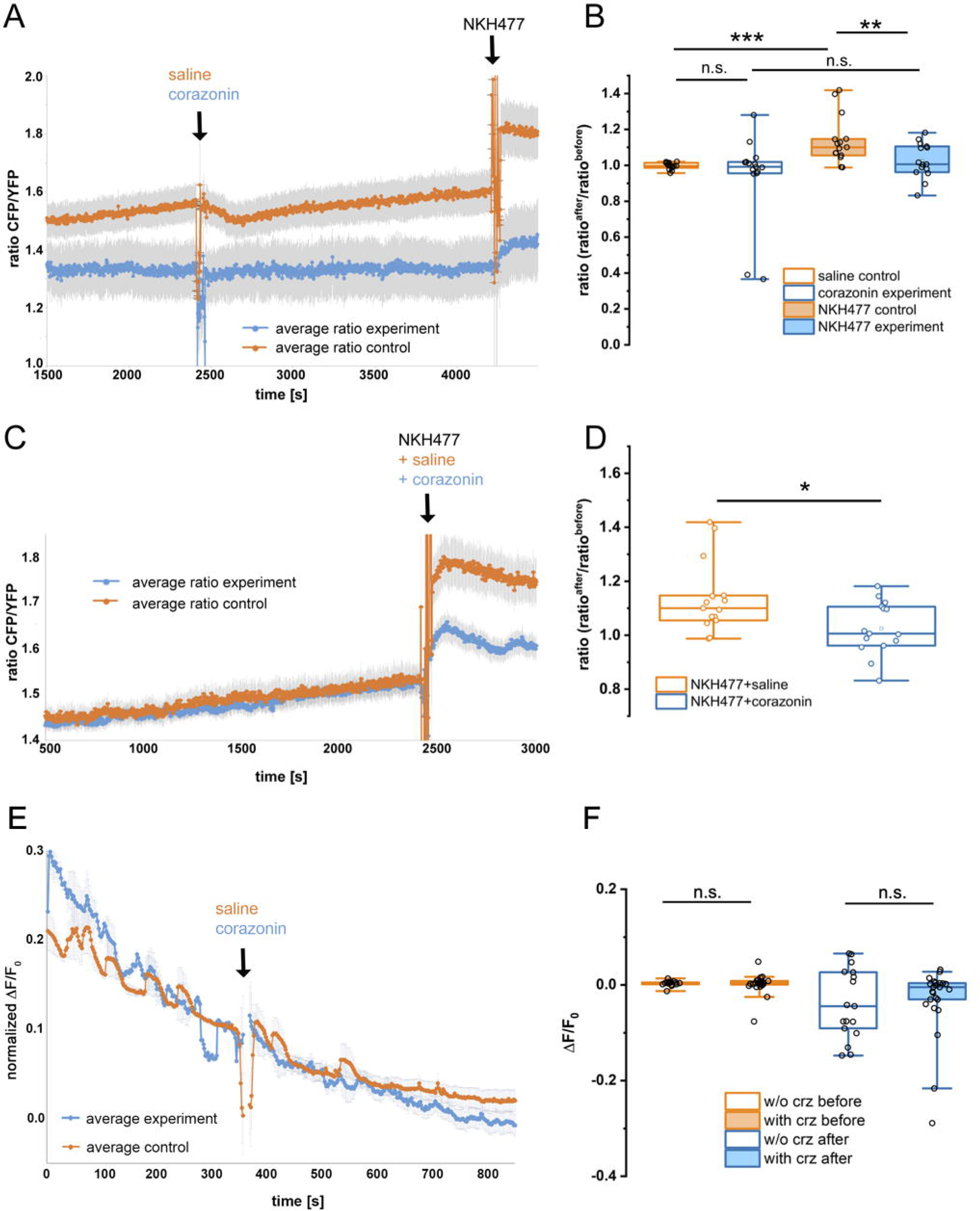
Crz inhibits cAMP levels in Va neurons. **(A, B)** Neither the application of 10 μM corazonin nor control saline (HL3.1 containing 0.1% DMSO) significantly altered the signal of the cAMP sensor Epac-camps. Under the experimental conditions, intracellular cAMP levels seemed to be rather low, as indicated by the strong and significant increase in the cAMP sensor signal after application of the membrane-permeable adenylyl cyclase activator NKH477 at the end of the control experiment. NKH477, however, failed to induce a significant rise in the cAMP sensor signal in the corazonin experimental group (n=15 cells, N=5 preparations for both experimental and control group). **(C, D)** Co-application of NKH477 with control saline induced a strong increase in cAMP signal. This NKH477-dependent increase was significantly diminished by co-application of corazonin (experimental group: n=23 cells, N = 5 preparations; control group: n=18 cells, N=5 experiments). **(E, F)** Application of 10 μM corazonin or control saline did not significantly affect the signal of the calcium sensor GCaMP6m (experimental group: n=25 cells, N=9 preparations; control group: n=17 cells, N=6 preparations. The decline of the signal in (E) represents photobleaching; the raw signal without baseline correction is shown. Grey bars in (A, C, E) represent s.e.m.

We next asked what the neuronal source of Crz is and whether its mode of action on Va neurons is systemic via the circulation or synaptic/paracrine within the abdominal neuromeres. To address this, we first examined the morphology of Crz and CAPA neurons. As described above, Crz is expressed in the DLPs, and in males, in abdominal ganglion interneurons **(S1 Figure)**. The DLPs do not have axonal projections descending into the VNC but they send axons to the corpora cardiaca (CC), anterior aorta and intestine (9, 46). Thus, Crz can be released into the hemolymph and is likely to act hormonally on Va neurons. The male-specific Crz interneurons in the abdominal neuromeres, on the other hand, could potentially interact with Va neurons via superimposed processes, at least in a paracrine fashion **(S8A Figure)**. However, the fact that *CrzR* expression is also detected in Va neurons of females **(S8B-C Figures)** suggests that the Crz-CAPA pathway is not restricted to males. In order to validate these assumptions, we utilized *trans*-Tango, an anterograde trans-synaptic labeling technique (47). *Crz-GAL4* driven *trans-*tango revealed expression of the post-synaptic marker across various regions of the brain and VNC **(S8D-E Figures)**. The absence of the post-synaptic signal in Va neurons of both males and females **(S8E Figure)** indicates that neither the DLPs nor the male-specific Crz interneurons are synaptically connected to the Va neurons. Thus, the DLPs, that are present in both sexes, appear to be the source of hormonal Crz, modulating the Va neurons.

### Release of Crz and CAPA peptides is altered under desiccation conditions

Next, we sought to determine whether Crz is released from the DLPs into the hemolymph during stress. We utilized Crz-immunoreactivity as a measure of peptide release; a decrease in Crz-immunoreactivity would indicate increased peptide release assuming there is no change in peptide production. The Crz immunoreactivity levels were quantified both in the large DLP neurons separately and also in all DLPs together, to see if there is a functional difference between these subpopulations **(Figure 4A)**. Interestingly, the Crz peptide level in the large DLP neurons was significantly reduced in desiccated and rewatered flies **(Figure 4A and B)** whereas the peptide level in all Crz neurons displayed a reduction in starved flies as well **(Figure 4A and S9A Figure)**. Since *Crz* transcription (a proxy for Crz peptide production) remains constant under osmotic **(Figure 4C)** and nutritional **(S9B Figure)** stresses, these results imply that there is stress-dependent release of Crz from the two subclasses of DLPs, albeit the cell-specific release varies depending on the type of stress. We also quantified *Capa* transcript levels following different stresses. Levels of *Capa* transcript are higher following desiccation **(Figure 4D)** as reported previously (26). Unlike its role during osmotic stress, the *Capa* transcript levels **(S9C Figure)** are unaltered following starvation (26) and refeeding suggesting that it is not involved in nutritional stress. Given that *Crz* knockdown led to increased chill-coma recovery time **(Figure 1B)**, we examined if *Crz* levels changed after chilling. Compared to normal flies, chilling and recovered flies demonstrated no change in *Crz* transcript levels **(Figure 4E)**, whereas comparatively, *Capa* transcript levels were significantly decreased in response to chilling and returned to normal levels when flies were allowed to recover at room temperature after chilling **(Figure 4F)**.

**Figure 4:**
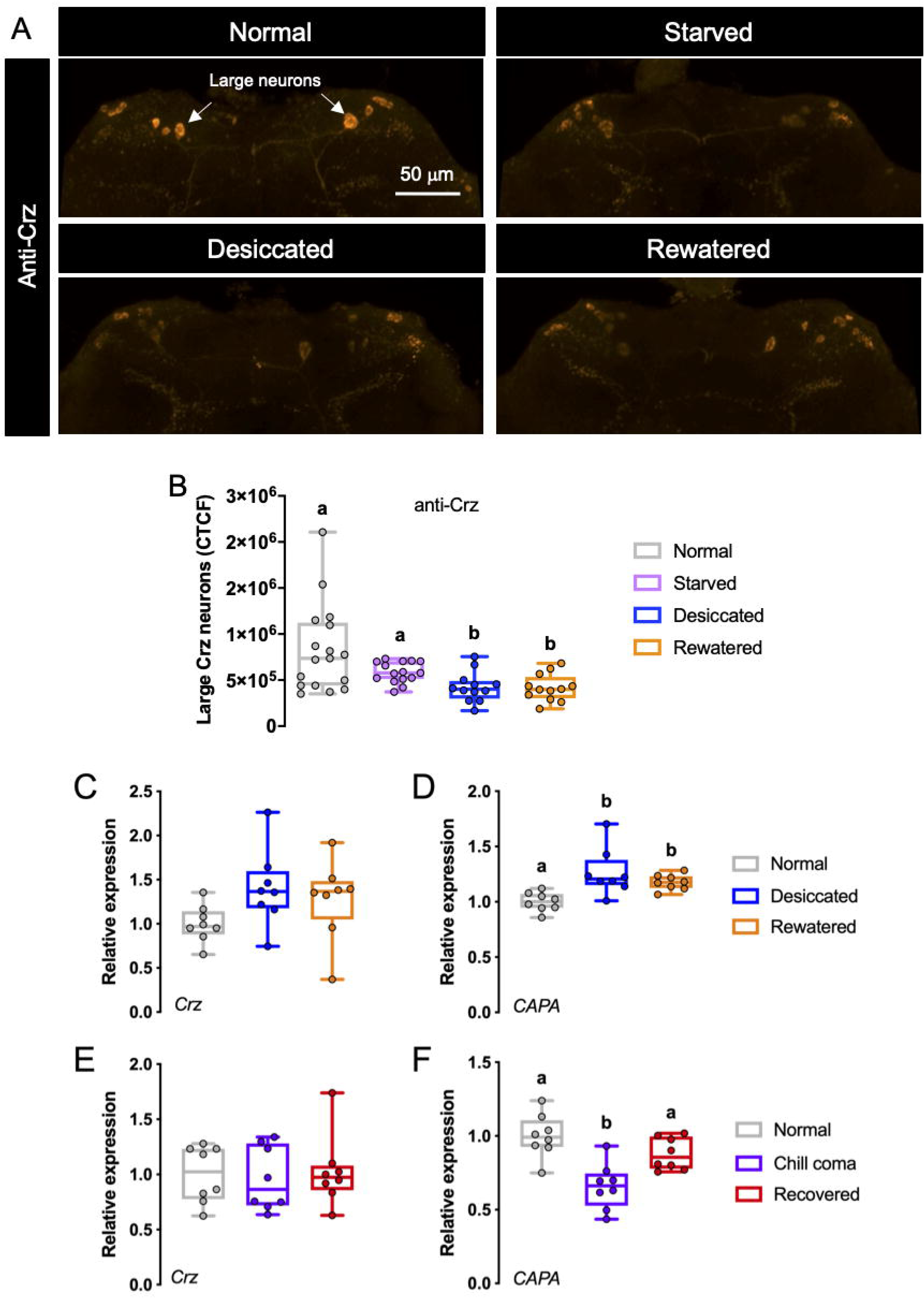
Crz peptide and transcript levels following nutritional and osmotic stresses. **(A, B)** Crz peptide levels in large neurosecretory cells (indicated by white arrows) are lower in desiccated and rewatered flies compared to flies raised under normal conditions (p < 0.01, assessed by One-way ANOVA). **(C)** *Crz* transcript levels are unaltered in desiccated and rewatered flies. **(D)** *CAPA* transcript levels are upregulated following desiccation and remain high after rewatering (p < 0.05, assessed by One-way ANOVA). **(E)** *Crz* transcript levels are unaltered following chill-coma. **(F)** *CAPA* transcript levels are downregulated following chill-coma and return to normal levels after recovery at room temperature (p < 0.05, assessed by One-way ANOVA).

### Crz signaling to Va neurons affects tolerance to osmotic stress

Since our imaging data indicates expression of *CrzR* in CAPA-producing Va neurons, and that Crz might inhibit CAPA release, we sought to examine the functional interaction between Crz signaling and Va neurons. To accomplish this, we first asked what role does CAPA play following its release into the hemolymph? CAPA was originally identified as a stimulator of fluid secretion in *Drosophila* (48) and since then it has been shown to stimulate secretion by Malpighian tubules of various insects, *ex vivo* (27, 49–53). However, the role of CAPA signaling has not been demonstrated *in vivo*. Thus, we monitored the excretion rate of flies in which CAPA neurons were optogenetically activated using ectopically expressed Channelrhodopsin-2, *ChR2-XXL*. Consistent with the *ex vivo* data, *CAPA>ChR2-XXL* flies exposed to light display increased defecation compared to genetically identical flies not exposed to light **(Figure 5A)**. This increased excretion could be mediated via actions on the gut and Malpighian tubule principal cells, which express the CAPA receptor (*CAPAR)* **(S10 Figure)** (54). In order to validate that Crz inhibits CAPA signaling, we again utilized optogenetics to simultaneously activate CAPA and Crz neurons. Activation of both the neuronal subsets abolishes the increased defecation seen after activation of CAPA neurons alone reiterating that Crz inhibits CAPA signaling **(Figure 5A)**. Moreover, if CAPA neuron activation results in increased defecation, then flies with increased CAPA signaling will retain less water and waste products. Indeed, activation of all CrzR-positive neurons (which include CAPA neurons) cause the flies to retain less water **(Figure 5B)** <u>whereas this decrease is not observed in</u> the no-GAL4 control flies **(S11A Figure)**. Taken together, our findings suggest that CAPA promotes excretion *in vivo* and Crz inhibits its release.

**Figure 5:**
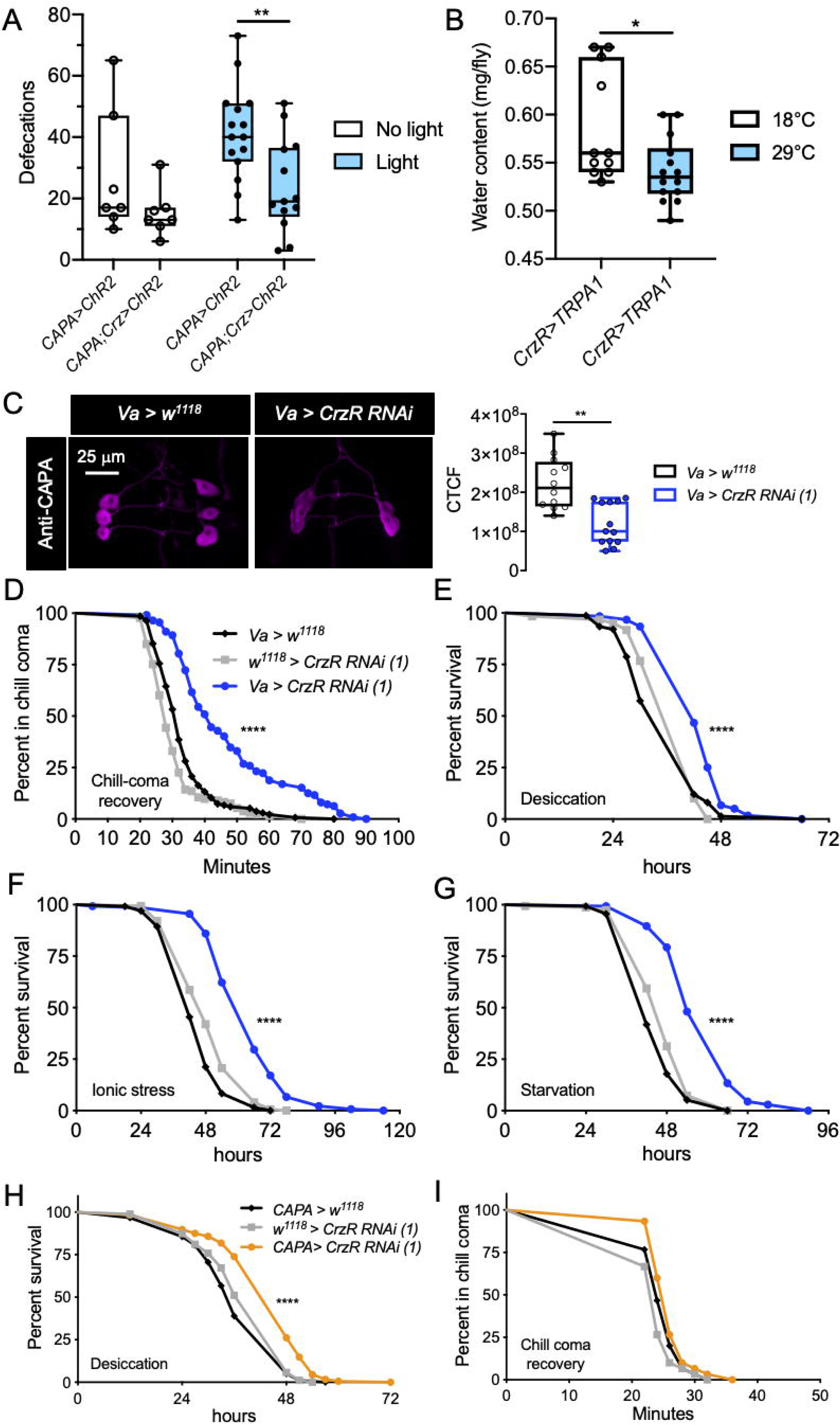
Functional interaction between Crz and CAPA signaling impacts water homeostasis and stress tolerance. **(A)** Optogenetic activation of CAPA neurons results in increased defecation and simultaneous activation of both CAPA and Crz neurons abolishes this increase (** p < 0.01 as assessed by unpaired *t*-test) **(B)** Thermogenetic TRPA1-driven activation of CrzR neurons in *CrzR > TRPA1* flies results in reduced water content (* p < 0.05 as assessed by unpaired *t*-test). **(C)** Knockdown of *CrzR* in Va neurons results in decreased CAPA levels, as measured using immunohistochemistry (** p < 0.01 as assessed by Mann-Whitney test). *Va-GAL4* driven *CrzR-RNAi(1)* results in **(D)** delayed recovery from chill-coma, and **(E)** increased survival under desiccation, **(F)** ionic and **(G)** starvation stress. *CAPA-GAL4* driven *CrzR-RNAi (1)* increases survival under **(H)** desiccation stress but has no impact on **(I)** chill-coma recovery. For **D-I**, data are presented as survival curves (**** p < 0.0001, as assessed by Log-rank (Mantel-Cox) test).

To further establish the interaction between Crz and CAPA signaling, we utilized independent GAL4 lines (*CrzR-GAL4*, *Va-GAL4, CAPA-GAL4*) to knock down the *CrzR* in various sets of neurons. Since both the *CrzR-RNAi* lines [*CrzR-RNAi (1)* and *(2)*] tested are equally effective at knocking down *CrzR*, we utilized *CrzR-RNAi (1)* (referred to as *CrzR-RNAi* from hereon) for all the experiments **(S11B Figure)**. The *CrzR-GAL4* drives broad expression in the CNS as described earlier, as well as in peripheral tissues such as the fat body. The *Va-GAL4* is stronger (based on the intensity of GFP expression) and drives expression in a broader set of neurons compared to *CAPA-GAL4* (26, 55). The *Va-GAL4* is expressed in several neurons in the brain and VNC in addition to the *CAPA* expressing neurons (**S12A-B Figures)**, whereas the *CAPA-GAL4* is only expressed in a pair of CAPA-negative brain neurons **(S12C Figure)** and the Va neurons in the VNC **(S12D Figure)**. Both the Va-GAL4 and CAPA-GAL4 have been used previously to manipulate gene activity in Va neurons (26). So using these three drivers, we asked whether knockdown of *CrzR* impacts phenotypes associated with CAPA signaling. Similar to that seen following global *Crz* knockdown, broad knockdown of *CrzR* with *CrzR-GAL4* also resulted in increased survival during desiccation **(S13A Figure)** and ionic stress **(S13B Figure)**. However, there was no effect on chill-coma recovery **(S13C Figure)**. These results further illustrate that disrupted Crz signaling influences tolerance to ionic and osmotic stresses.

Next, we wanted to examine the effects of knocking down *CrzR* in smaller populations of neurons including those which produce CAPA. If Crz inhibits CAPA release, knockdown of *CrzR* in Va neurons should relieve this inhibition and result in increased CAPA release. Consistent with this prediction, *Va-GAL4* driven *CrzR-RNAi* indeed results in decreased CAPA peptide levels in Va neurons **(Figure 5C)**. Since *Capa* transcript levels are unaltered in *CrzR* knockdown flies **(S11C Figure)**, these results indicate that *CrzR* knockdown causes increased CAPA release. Additionally, if knockdown of *CrzR* in Va neurons affects CAPA release, then phenotypes associated with CAPA signaling should consequently be impacted. Indeed, *Va-GAL4* driven *CrzR-RNAi* results in delayed recovery from chill-coma **(Figure 5D)** and increased survival during desiccation **(Figure 5E)** and ionic stress **(Figure 5F)**, consistent with the role of CAPA as an osmo- and ionoregulatory peptide. In addition, we observed increased survival under starvation **(Figure 5G)**, possibly due to effects on food intake as a result of *CrzR* knockdown in non-CAPA expressing neurons **(S14 Figure)**. Hence, we knocked down the *CrzR* more selectively using *CAPA-GAL4*. The *CAPA-GAL4* driven *CrzR-RNAi* resulted in increased survival under desiccation **(Figure 5G)**. However, it did not have any impact on chill-coma recovery **(Figure 5H)**. Similar results were also obtained by knocking down *CrzR* in CAPA neurons using an independent *CrzR-RNAi* construct *CrzR-RNAi (2*). The lack of effect on chill-coma recovery following *CrzR* knockdown with *CAPA-GAL4* could be due to it being a very weak driver. Taken together, our data indicate that Crz acts on Va neurons to inhibit the release of CAPA, and thereby affecting tolerance to osmotic and ionic stresses.

## Discussion

We have found here that a peptidergic neuroendocrine pathway in *Drosophila*, known to restore nutrient deficiency (utilizing Crz), integrates a further peptidergic component (CAPA) to maintain osmotic and ionic homeostasis (**Figure 6** **and Table S1)**. The Crz-CAPA signaling thereby also influences tolerance to osmotic and cold stress. An earlier study suggested that Crz is released during nutritional stress to mobilize energy stores from the fat body to fuel food search behavior (14). Furthermore, that study suggests that increased Crz signaling compromises resistance to starvation, desiccation and oxidative stress (14). We confirm these findings here, and also find that Crz inhibits a set of *CrzR* expressing Va neurons in the abdominal ganglia that produce CAPA peptides, which when released *in vivo* through optogenetic or thermogenetic control, leads to increased excretion and decreased whole body water content, respectively. Thus, the two peptidergic systems act together to maintain both energy and ion/water homeostasis.

**Figure 6:**
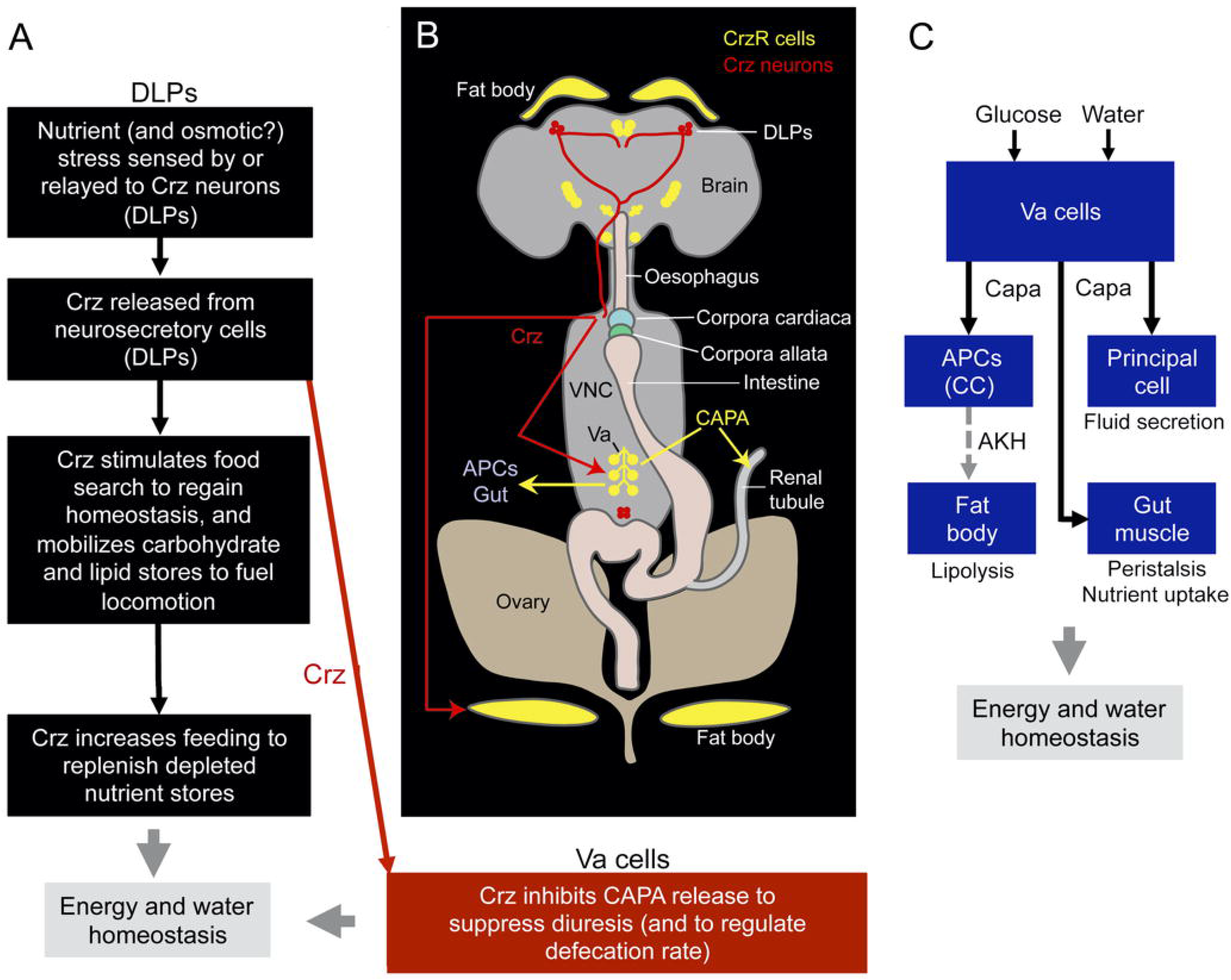
Crz signaling modulates energy and osmotic homeostasis. **(A)** A proposed model showing the mode of action of Crz in modulating response to starvation leading to increased food search and food ingestion (black arrows). If the fly is exposed to starvation, especially dry starvation, there is an acute response to preserve water and release of the diuretic peptides (CAPA1 and 2) is inhibited (red arrow). When the starved or desiccating fly monitors low nutrient levels (later than the response to osmotic stress) the Crz release ensues and the peptide acts to mobilize energy stores to fuel food search. After feeding and restoration of energy homeostasis, Crz inhibition of Va neurons ceases and CAPA release can restore postfeeding water homeostasis. **(B)** A schematic showing the location of Crz (red) and CrzR-expressing neurons/cells (yellow), as well as a model proposing how Crz modulates nutritional and osmotic stresses. Crz is released from dorsal lateral peptidergic neurons (DLPs) into the hemolymph and activates its receptor on the fat body and Va neurons. Crz modulates the release of CAPA from Va neurons, which in turn affects chill-coma recovery and desiccation tolerance via its effect on the renal tubules. CAPA is also acting on AKH-producing cells (APCs) and the gut, as shown in (61). Note that *CrzR* is also expressed in the antennal lobes; however, these have been excluded from the figure for purposes of clarity. (C) Model redrawn from (61) depicting roles of Va neurons in regulation of energy and water homeostasis. The activity in Va neurons increases in response to feeding and drinking (glucose and water). CAPA is released to act on APCs in corpora cardiaca (CC), renal tubules (principal cells) and intestinal muscle. CAPA inhibits AKH release from APCs (dashed line). Activation of Va cells leads to decreased lipolysis as well as increased fluid secretion, gut peristalsis and nutrient uptake. The schemes are based on findings in this study and those of previous reports (14, 17, 25, 26, 46, 61, 63).

The *Capa* gene-derived neuropeptides (CAPA1 and CAPA2) are well established as osmoregulatory factors that act on the Malpighian tubule principal cells (see 56, 57) and perhaps the hindgut, as indicated herein by our receptor expression data. Previous *ex vivo* studies have largely showed that CAPA neuropeptides act as diuretic hormones in *Drosophila* and other dipterans (31, 58) while anti-diuretic actions have also been reported (27, 51, 59, 60). Our *in vivo* findings herein are more aligned with the observations supporting a diuretic role of CAPA peptides Furthermore, we show that the CAPA-producing Va neurons are downstream of Crz signaling and we propose that under adverse conditions when flies are exposed to dry starvation (desiccation without food) the two signaling systems act in tandem to restore homeostasis. When the fly experiences starvation and nutrients diminish in the fly, nutrient sensors record this deficiency, which triggers release of hormones that act to restore metabolic homeostasis. As indicated above, one such hormone is Crz, known to provide energy for food search and induce feeding (14, 16). The released Crz also inhibits CAPA release from Va cells during the nutrient shortage and results in diminished excretion (and defecation) during starvation and food search. Additionally, during the revision of this manuscript, a recent preprint (61) reported that CAPA from Va cells also acts on AKH producing cells in the corpora cardiaca to inhibit AKH release and thereby decreasing lipolysis in adipocytes. Thus, when Crz inhibits the Va neurons, AKH signaling is likely to be elevated and results in increased energy mobilization from the fat body, further facilitating food search (Fig. 6). In insects, increased food intake yields water (see 62), which requires post-feeding diuresis to be activated to restore water homeostasis. Therefore, the inhibition of CAPA release from Va cells needs to be lifted after successful food search and nutrient intake. This suggests that the Crz action occurs during the starvation/fasting, and thereafter postprandial CAPA action sets in to restore nutrient and water homeostasis. Unfortunately, we cannot provide direct evidence for this sequence of events since there is no method sensitive enough to monitor the timing of changes in hemolymph levels of Crz and CAPA in small organisms such as *Drosophila*. Indirect measurements, like monitoring levels of peptide-immunolabel in neurons of interest are not necessarily accurate, since these levels reflect the “balance” between peptide release and production and therefore not useful for resolving onset and duration of release with accuracy.

Our evidence for the Crz action on CAPA-producing Va neurons is based on the expression of *CrzR* in these cells, as well as functional imaging data which shows that Crz inhibits cAMP production in Va neurons (but has no impact on intracellular Ca^2+^ levels). This led us to investigate the effects of manipulating Crz signaling and targeted *CrzR* knockdown in Va neurons on aspects of water and ion homeostasis and cold tolerance, which have been shown earlier to be affected by CAPA signaling (26, 27). We observe here that global Crz or CrzR knockdown leads to delayed recovery from chill-coma (i.e. jeopardizing cold tolerance) and increased tolerance to ionic stress. Furthermore, our experiments that target knockdown of CrzR to Va neurons also affected chill-coma recovery, and tolerance to starvation, desiccation, and ionic stresses. Injecting flies *in vivo* with Crz in the present study also resulted in effects on chill-coma recovery and survival during desiccation, further strengthening our model in which Crz modulates release of CAPA. However, the effect of CAPA injections on chill-coma recovery shown previously (27) are not directly compatible with our findings. This may be due to the difficulty in comparing the effects obtained from RNAi manipulations (chronic) and peptide injections (acute). Furthermore, Crz not only affects cold tolerance via its actions on CAPA signaling, it also regulates levels of trehalose, a cryoprotectant, via actions on the fat body (14). Thus, Crz signaling could modulate cold tolerance via two independent hormonal pathways.

Previous work in *Drosophila* showed that not only does global *Crz* knockdown result in increased resistance to starvation, desiccation and oxidative stress, but also found that *CrzR* knockdown in the fat body (and salivary glands) led to these same phenotypes (9, 14). These findings suggest that Crz signaling to the fat body accounts for some of the stress tolerance phenotypes seen following *Crz* knockdown. Thus, it is possible that the effects observed on desiccation survival following Crz peptide injections in the present study are partly confounded by the actions of this peptide on the fat body and other tissues expressing the CrzR.

Earlier data show that after *CrzR* knockdown in the fat body, glucose, trehalose and glycogen levels are elevated in starved flies, but not in normally fed flies (14). Moreover, starved, but not normally fed flies with *Crz* knockdown, display increased triacyl glycerides (9) and *Crz* transcript is upregulated in starved flies with *CrzR* knockdown in the fat body (14). This further emphasizes that systemic Crz signaling is critical under nutritional stress. Our data on Crz immunolabeling intensity corroborate these earlier findings and suggest that Crz is released to restore metabolic homeostasis by mobilizing energy stores from the fat body to fuel food search behavior. After restoring nutritional and osmotic stress the Crz signaling decreases and thus the proposed inhibition of CAPA release from Va neurons is lifted.

We did not address the question as to how the Crz neurons sense nutrient deficiency or whether they can detect changes in osmolarity (but see below). However, it is known that Crz-producing DLPs express an aquaporin, Drip, and the carbohydrate-sensing gustatory receptors, Gr43a and Gr64a (63–65). A subset of the Crz neurons also express a glucose transporter (Glut1) that is involved in glucose-sensing (17). Thus, imbalances in internal nutritional and maybe even osmotic status could either be sensed cell autonomously by the Crz neurons or indirectly by signals relayed to them via other pathways.

A few other findings might support roles of Crz in ameliorating nutrient stress. We found here that Crz neurons innervate the antennal lobe (AL), and that the *CrzR* is strongly expressed in local interneurons of the AL. Possibly, Crz modulates odor sensitivity in hungry flies to increase food search, similar to peptides like short neuropeptide F (sNPF), tachykinin and SIFamide (66–68). Another peptide hormone, adipokinetic hormone (AKH), has been shown to be critical in initiating locomotor activity and food search in food deprived flies (13, 15, 69) and AKH also affects sensitivity of gustatory neurons to glucose (70, 71). The effect of AKH on increasing locomotor activity is evident only after 36h of starvation (69) and may correlate with the proposed action of Crz during starvation (see 14). Possibly Crz acts in concert with AKH to allocate fuel during metabolic stress (69). Since we found that the *CrzR* is not expressed in AKH-producing cells, it is likely that these two peptides act in parallel rather than in the same circuit/pathway. However, the DLPs also produce sNPF (9) and this peptide is known to act on the AKH producing cells and thereby modulate glucose homeostasis and possibly, sensitivity of gustatory neurons (17, 19, 70). Thus, the DLPs may act systemically with Crz and by paracrine signaling in the corpora cardiaca to act on AKH cells. Thereby the Crz and AKH systems could be linked by sNPF. In addition, while this manuscript was under revision, a preprint (61) was posted that added an interesting angle to the role of AKH signaling. That study revealed that the AKH producing cells (APCs) express the CAPA receptor and that CAPA acts on APCs to decrease AKH release, which diminishes lipolysis in adipocytes. Thus, in the fed fly, CAPA not only induces diuresis, it also diminishes energy mobilization (61). The same study also showed that the Va cells in flies are active after sucrose feeding or drinking, and that CAPA triggers nutrient uptake and peristalsis in the intestine (see **Fig. 6C**). This interaction between CAPA-AKH signaling can also explain the direction of the phenotypes seen following CrzR knockdown in Va neurons. For instance, knockdown of CrzR in Va neurons results in increased CAPA release and one would predict decreased desiccation survival if CAPA is exclusively promoting excretion. However, increased CAPA signaling could also result in reduced AKH signaling (61) that in turn promotes starvation survival (15). Thus, the direction of the phenotypes we observe following CrzR knockdown in Va neurons can be explained by CAPA actions on excretion and AKH-release from APCs. Lastly, we also do not rule out the possibility that other neuropeptides possibly coexpressed with CAPA (37) could also contribute to the observed phenotypes.

In conclusion, we suggest that Crz regulates acute metabolic stress-associated physiology and behavior via the fat body to ensure nutrient allocation to power food search and feeding during prolonged starvation. During food search and feeding, excretion is blocked by Crz acting on the Va cells to inhibit CAPA release. Following food intake and ensuing need for diuresis, Crz signaling ceases and CAPA can be released to ensure restoration of water and ion homeostasis. Taken together, our findings and those of previous studies indicate that Crz acts on multiple neuronal and peripheral targets to coordinate and sustain water, ion and metabolic homeostasis. It might even be possible that an ancient role of the common ancestor of Crz and AKH signaling systems was to modulate stress-associated physiology and that these paralogous signaling systems have sub-functionalized and neo-functionalized over evolution (22, 72).

## Materials and methods

### Fly lines and husbandry

*Drosophila melanogaster* strains used in this study are listed in **Table 1**. Flies were backcrossed into the same genetic background (*w*^*1118*^) for 7 generations, with exception of the stocks used for imaging and defecation assays. All stocks were stored at 18°C under normal photoperiod (12 hours light: 12 hours dark; 12L:12D) on a standard cornmeal/molasses/yeast diet. Unless indicated otherwise, experimental flies were reared under non-crowded conditions and maintained under normal photoperiod at 25°C on enriched medium containing 100 g/L sucrose, 50 g/L yeast, 12 g/L agar, 3 ml/L propionic acid and 3 g/L nipagin. Adult males 6-7 days old post-eclosion were used unless mentioned otherwise.

**Table 1:**
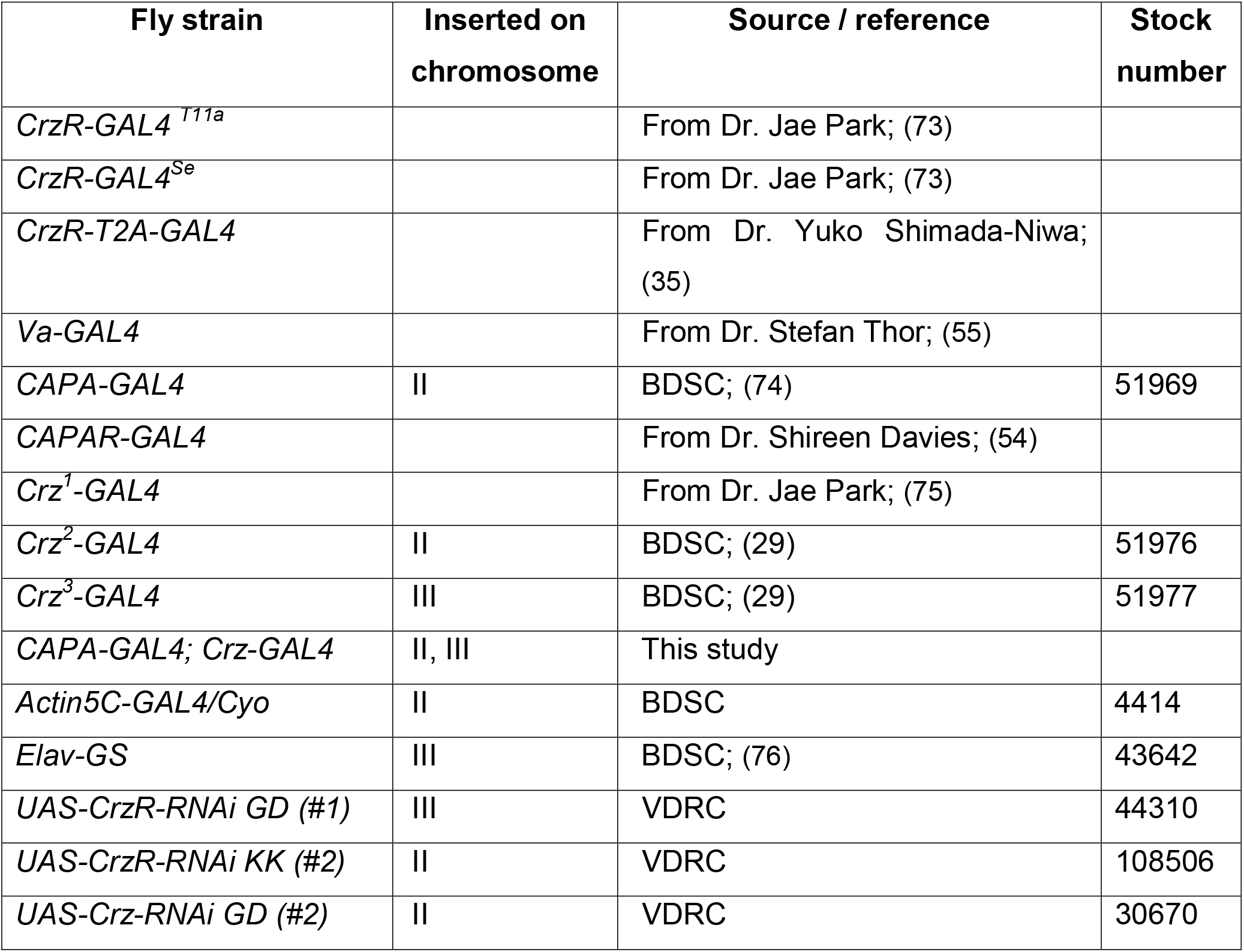
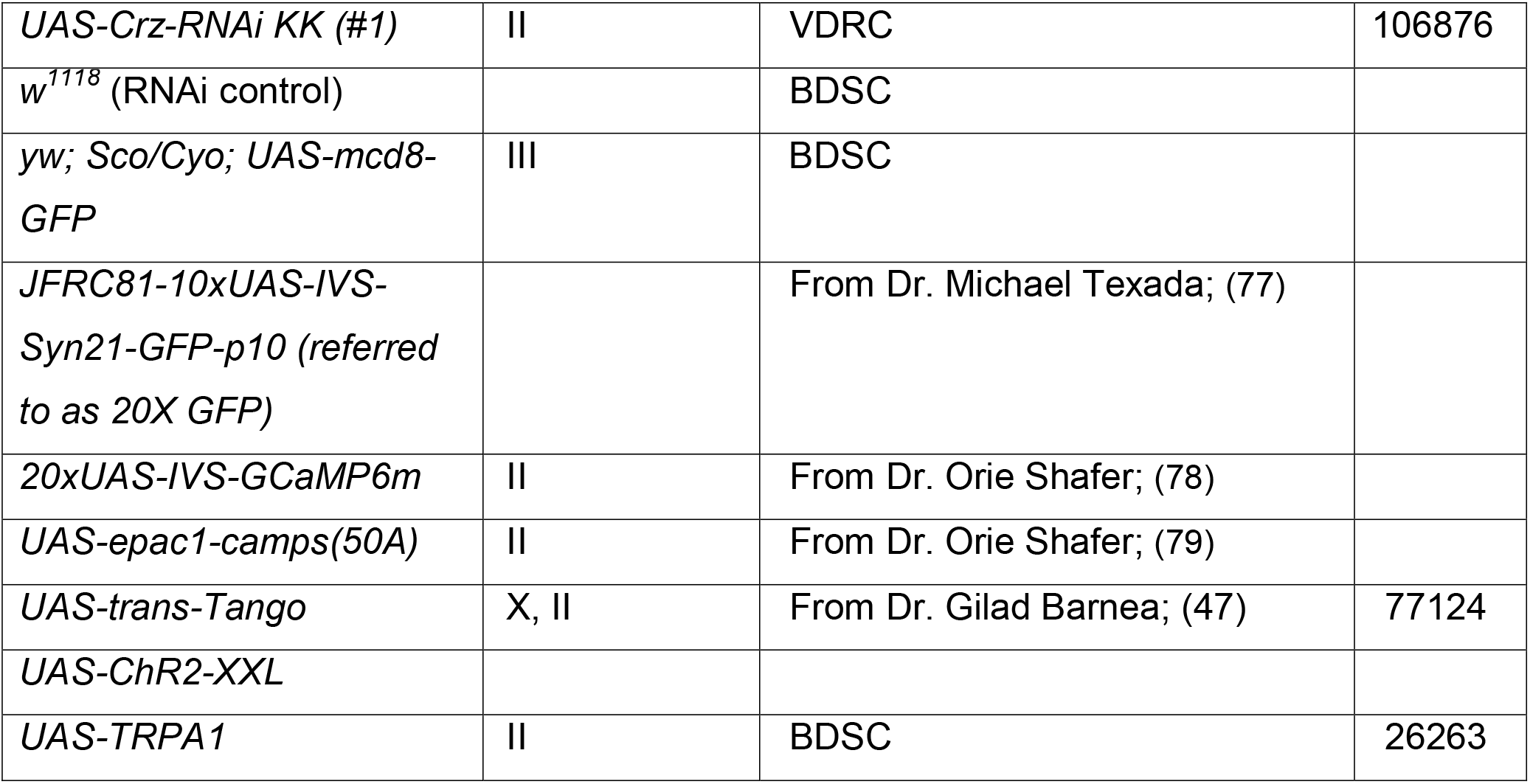
Fly strains used in this study

### Immunohistochemistry and imaging

Immunohistochemistry for *D. melanogaster* adult CNS and gut was performed as described earlier (34). Briefly, tissues were dissected in phosphate buffered saline (PBS), fixed in 5% ice-cold paraformaldehyde (3.5-4 hours), washed in PBS and incubated in primary antibodies (**Table 2**) diluted in PBS with 0.5% Triton X (PBST) for 48 hours at 4°C. Samples were then washed with PBST and incubated in secondary antibodies (**Table 2**) diluted in PBST for 48 hours at 4°C. Finally, samples were washed with PBST and then PBS before being mounted in 80% glycerol. Zeiss LSM 780 confocal microscope (Jena, Germany) was used to image all the samples.

**Table 2:** Antibodies used for immunohistochemistry

| Antibody | Immunogen | Source / reference | Dilution |
| --- | --- | --- | --- |
| ***Primary antibodies*** |  |  |  |
| Rabbit anti-PVK-2<br>(referred to as anti-PRXa in Figure S6C) | *Periplaneta americana*<br>CAPA-PVK-2 | Dr. Reinhard Predel<br>(81) | 1:4000 |
| Rabbit anti-CAPA-2<br>(referred to as anti-CAPA in text and other figures) | *Rhodnius prolixus* CAPA-2 | Dr. Ian Orchard (27) | 1:1000 |
| Rabbit anti-Crz | *Drosophila* Crz | Dr. Jan Veenstra<br>(82) | 1:4000 |
| Rabbit anti-DILP2 | *Drosophila* DILP2 | Dr. Jan Veenstra<br>(83) | 1:2000 |
| Rabbit anti-AKH | *Drosophila* AKH | Dr. Mark Brown (84) | 1:1000 |
| Rabbit anti-FMRFamide | FMRFamide | Dr. Cornelis<br>Grimmelikhuijzen<br>(85) | 1:4000 |
| Rabbit anti-PDH/PDF | Crab *Uca pugilator*<br>pigment dispersing<br>hormone | Dr. Heinrich<br>Dircksen (86) | 1:4000 |
| Rabbit anti-LK | *Leucophaea maderae* kinin<br>I | (87) | 1:2000 |
| Rabbit anti-DH44 | *Drosophila* DH44 | (88) | 1:1000 |
| Rabbit anti-DH31 | *Drosophila* DH31 | (89) | 1:1000 |
| Rat anti-HA | Monoclonal Clone 3F10 | Sigma | 1:100 |
| Mouse anti-GFP | Jelly fish GFP | Invitrogen | 1:1000 |
| Chicken anti-GFP | Jellyfish GFP | Invitrogen | 1:1000 |
| Goat anti-GFP | Jellyfish GFP | Rockland | 1:1000 |
| Mouse anti-nc82 | *Drosophila* bruchpilot | DSHB | 1:20 |
| ***Secondary antibodies*** |  |  |  |
| Goat anti-mouse<br>Alexa 488 | - | Invitrogen | 1:1000 |
| Goat anti-rabbit Alexa<br>546 | - | Invitrogen | 1:1000 |
| Goat anti-chicken<br>Alexa Fluor 488 | - | Life Technologies | 1:1000 |
| Goat anti-mouse<br>Alexa Fluor 546 | - | Life Technologies | 1:1000 |
| Donkey anti-rabbit<br>Alexa Fluor 647 | - | Life Technologies | 1:1000 |
| Goat anti-rabbit<br>Cyanine5 | - | Life Technologies | 1:500 |
| Donkey anti-rat<br>Alexa Fluor 555 | - | Southern Biotech | 1:1000 |
| Donkey anti-goat<br>Alexa Fluor 488 | - | Invitrogen | 1:1000 |
| ***Other fluorophores*** |  |  |  |
| Rhodamine-phalloidin | - | Invitrogen | 1:1000 |

To quantify peptide levels in flies exposed to various stressors, adult males were transferred to either an empty vial (desiccation) or a vial containing aqueous 0.5% agar (starvation) or artificial diet (normal food) and incubated for 18 hours. In addition, one set of flies were desiccated for 15 hours and then transferred to a vial containing 0.5% agar (re-watered) for 3 hours. These flies were then processed for immunohistochemistry as described above. Cell fluorescence was quantified as described previously (34). Note that the anti-CAPA antibody used to quantify CAPA peptide levels cross-reacts with other PRXamide-related peptides. Hence, this approach measures the levels of all PRXamide peptides that are coexpressed in a given neuron.

For *trans*-Tango experiments, flies were raised at 18°C and 3 weeks old adult males and females were used for immunohistochemistry.

Confocal images were processed with Fiji (80) for projection of z-stacks, contrast and brightness, and calculation of immunofluorescence levels. Further adjustments (cropping and brightness) were made in Microsoft Powerpoint.

### Stress resistance assays

Flies were assayed for survival under desiccation, starvation and ionic stress by being kept in empty vials, vials containing 0.5% aqueous agarose (A2929, Sigma-Aldrich), and vials containing artificial food supplemented with 4% NaCl, respectively. Flies were kept in vials and their survival recorded every 3 to 6 hours until all the flies were dead. The vials were placed in incubators at 25°C under normal photoperiod conditions (12L:12D). For chill-coma recovery of transgenic flies, flies were incubated at 0°C for 4 hours and then transferred to room temperature (24°C) to monitor their recovery time. At least three biological replicates and two technical replicates (10-15 flies per technical replicate) for each biological replicate were performed for each experiment.

For experiments using the GeneSwitch system (30), flies were reared on regular food. Following 3-4 days post-eclosion, half the males were transferred to regular food and the other half transferred to food containing 200μM RU486, and incubated for another 4 days prior to using them for the ionic stress assay.

### Peptide injection experiments

*Drosophila w*^*1118*^ males 7-8 days post-eclosion were used for these experiments. These flies were reared and maintained at room temperature (~23°C).

#### Peptide Injections

*D. melanogaster* Crz (pQTFQYSRGWTN-NH_2_) was custom synthesized at >95% purity by Genscript (Piscataway, NJ, USA); a non-amidated adipokinetic hormone/corazonin-related peptide (naACP) from *Aedes aegypti* (pQVTFSRDWNA), as previously described (90), was custom synthesized at >90% purity by Pepmic Co. (Suzhou, Jiangsu, China). Peptides were initially solubilized in nuclease-free deionized water or DMSO to a stock concentration of 10^−3^ M and then each peptide was diluted in *Drosophila* saline injection buffer (NaCl 117.5 mM, KCl 20 mM, CaCl_2_ 2 mM, MgCl_2_ 8.5 mM, NaHCO_3_ 10.2 mM, NaH_2_PO_4_ 4.3 mM, HEPES 15 mM, glucose 20 mM, pH 6.7) as described previously (26) and supplemented with 0.1% (w/v) Fast Green FCF, which was used to visually confirm successful injections. Peptide injection solutions were prepared to achieve final concentrations in the hemolymph of either 10^−6^ or 10^−9^ M and based on the *Drosophila* hemolymph volume of 80 nL (91). Peptide injections took place at room temperature under light CO_2_ anesthesia, with injections directed to the left mesopleuron using a micromanipulator-controlled Drummond Nanoject Injector set to 18.4 nL per injection. Control flies received only *Drosophila* saline injection buffer containing 0.1% Fast Green FCF.

#### Chill-Coma Recovery

Flies were taken from the *Drosophila* vials and lightly anesthetized by CO_2_. Once immobilized, male flies were isolated and placed into a white plastic weighing dish that was held over ice. For each treatment, 9-15 males were selected per treatment. After injections, flies were transferred individually into 7.5 mL glass vials that were then submerged in a 0°C ice-water slurry for 1 hour. Afterwards, vials were removed from the slurry and gently dried. Flies were left to recover at room temperature without being disturbed for the duration of the experiment. Recovery was recorded by measuring the time when a fly stands on all legs. Experiments were repeated in at least two independent biological replicates.

#### Desiccation

As above, male flies were isolated and placed over ice within a white plastic weighing dish. For each treatment, 11-15 males were used. After injection with saline injection buffer alone or buffer containing Crz, all flies in that treatment group were transferred *en masse* into empty *Drosophila* vials without any food or water. Survival under this desiccation treatment was monitored at regular intervals. This experiment was repeated in three replicates.

### Calcium and cAMP imaging of Va neurons

The whole CNS of feeding 3^rd^ instar larvae expressing the calcium sensor GCaMP6m (Chen, Wardill et al. 2013) or the cAMP sensor Epac1-camps (Shafer, Kim et al. 2008) were dissected in HL3.1 saline ((92), pH 7.2-7.4) and attached at the bottom of a plastic Petri dish lid (35×10 mm; Greiner Bio-One International GmbH, Austria) filled with HL3.1. The CNS was allowed to settle for 10 min and then imaged on a widefield fluorescent imaging setup (for calcium imaging: Zeiss AxioExaminer D1, equipped with a W “Plan-Apochromat” x20/1.0 and a Chroma-ET GFP filter, a pco.edge 4.2 sCMOS camera and a SPECTRA-4 light engine; for cAMP imaging: Zeiss Axioskop FS2, equipped with a ZeissPhotometrics DualView2 with Chroma ET-CFP and ET-YFP emission filters and a dualband CFP/YFP dichroic mirror, a CoolSnap CCD camera and a VisiChrome monochromator; both systems from Visitron, Puchheim, Germany).

For calcium imaging, an excitation wavelength of 475 nm and an exposure time of 180 ms at 2x binning was used. Imaging was performed for 15 min at 1 Hz. After 350 seconds, 10 μM (end concentration) synthetic Crz dissolved in HL3.1 containing 0.1% DMSO or HL3.1 plus 0.1% DMSO (control) was added. For data analysis, background was subtracted and the change in fluorescence intensity was calculated as *ΔF*/*F*_*0*_ = (*F*_*n*_-*F*_0_)/*F*_*0*_ were *F*_*n*_ is the fluorescence at time point *n* and *F*_*0*_ is the mean baseline fluorescence value of the 30 s before peptide/control application. For cAMP imaging, an excitation wavelenght of 434/17 nm and an exposure time of 800 ms at 4x binning was used. Imaging was performed for 80 minutes at 1 Hz. In the first set of experiments, 10 μM synthetic Crz dissolved in HL3.1 containing 0.1% DMSO or HL3.1 plus 0.1% DMSO (control) was added after 40 min. At 75 minutes, 100 μM of the membrane-permeable adenylyl cyclase activator NKH477 (Merck Millipore) was added. In the second set of experiments, 100 μM NKH477 in 0.1% DMSO alone or in combination with 10 μM Crz was added. For data analysis, backgrounds were subtracted and the CFP/YFP ratio was calculated for each time point with a YFP signal corrected for spill-over.

### Quantitative PCR

#### CrzR-RNAi Knockdown Efficiency

Total RNA was isolated from whole male flies using Quick-RNA MiniPrep (Zymo Research) from four independent biological replicates with 8-15 flies in each replicate. The RNA concentration was determined using NanoDrop 2000 spectrophotometer (Thermo Scientific). RNA was diluted to ensure similar amount across treatments, and used to synthesize cDNA with random hexamer primers (Thermo Scientific) and RevertAid reverse transcriptase (Thermo Scientific). The cDNA was then diluted and used as template for qPCR, which was performed with a StepOnePlus instrument (Applied Biosystem, USA) and SensiFAST SYBR Hi-ROX Kit (Bioline) according to the manufacturer’s instructions. The mRNA levels were normalized to rp49 levels in the same samples. Relative expression values were determined by the 2^−ΔΔCT^ method (93). See **Table S1** for the primers used for qPCR.

#### Crz and Capa Transcript Levels Following Stress

To examine if stress exposure, including desiccation, starvation and chill coma, modified levels of *Crz* and *Capa* transcripts, *w*^*1118*^ male flies 6-7 days old were isolated into fresh vials containing enriched medium (with 20-25 flies/vial).

For desiccation experiments, two out of three vials of flies were transferred to empty vials for 18 hours. After 18 hours of desiccation, flies from one of the empty vials were then transferred into a fresh vial containing moistened compacted Kimwipes (4 wipes with 4mL H_2_O) for 4 hours. The remaining two vials, including control flies left in enriched medium for the duration of the experiment and flies desiccated for 18 hours, were sacrificed by plunging in liquid nitrogen and then frozen whole flies were stored at −80°C. Following the 4-hour recovery on moistened Kimwipes of the last vial of flies, the same procedure was done and these flies were sacrificed and stored in −80°C freezer.

For starvation experiments, two out of three vials of flies were transferred to vials containing moistened and compacted Kimwipes for 18 hours. After 18 hours of starvation, one of the two vials of flies with moistened Kimwipes was then transferred into a fresh vial containing enriched medium for 4 hours. The remaining two vials, including control flies left in enriched medium for the duration of the experiment and flies that had been starved for 18 hours, were immediately sacrificed as described above and stored in −80°C freezer. Following the 4-hour feeding recovery on enriched medium of the last vial of flies, they were sacrificed and stored in −80°C freezer.

For chill coma experiments, two out of three vials of flies were transferred to an ice water slurry and maintained at 0°C for 18 hours. After 18 hours of chill-coma, flies from one of the two vials chilled for 18 hours were transferred into a fresh vial containing enriched medium for 4 hours at room temperature. The remaining two vials, including control flies left in enriched medium for the duration of the experiment and flies that had been subjected to chill-coma for 18 hours, were immediately sacrificed as described above and stored in −80°C freezer. Following the 4-hour room temperature recovery of the last vial of flies, they were sacrificed and stored at −80°C. All the above experiments were repeated in three to four biological replicates.

Total RNA was isolated using the Monarch Total RNA Miniprep Kit (New England Biolabs, Whitby, ON) following manufacturer guidelines, which included gDNA removal. Purified total RNA was quantified on a Synergy 2 Multimode Microplate Reader using a Take3 micro-volume plate (BioTek, Winooski, VT). For cDNA synthesis, 300ng total RNA was used as template to prepare first-strand cDNA using the iScript Reverse Transcription Supermix for RT-qPCR (Bio-Rad, Mississauga, ON) in a total reaction volume of 10 μL. Following the cDNA synthesis reaction, cDNA was diluted 20-fold by adding nuclease-free H_2_O (Wisent, St. Bruno, QC, Canada) and samples were then stored at −20°C until ready for analysis using qPCR.

Transcript abundance of *Crz* and *Capa* in whole fly cDNA samples was quantified using PowerUp SYBR Green Master Mix (Applied Biosystems, Carlsbad, CA) using fast cycling mode on a StepOnePlus Real-Time PCR System (Applied Biosystems, Carlsbad, CA). Cycling conditions involved initial uracil-DNA glycosylase (UDG) at 50°C for 2 minutes, Dual-Lock DNA polymerase activation at 95°C for 2 minutes, followed by 40 cycles of denaturing at 95°C for 3 seconds and combined annealing extension step at 60°C for 30 seconds.

Following the amplification, reactions were assessed by melt curve analysis and products were also visualized by gel electrophoresis and sequence specificity confirmed by Sanger sequencing. Relative expression levels were determined using the 2^−ΔΔCT^ method (93) and normalized to transcript abundance of *rp49* used as a reference gene. Primers were designed over exon-exon boundaries when possible, or at minimum were localized on different exons to exclude possible gDNA amplification. See **Table 3** for the primers used for qPCR.

**Table 3:**
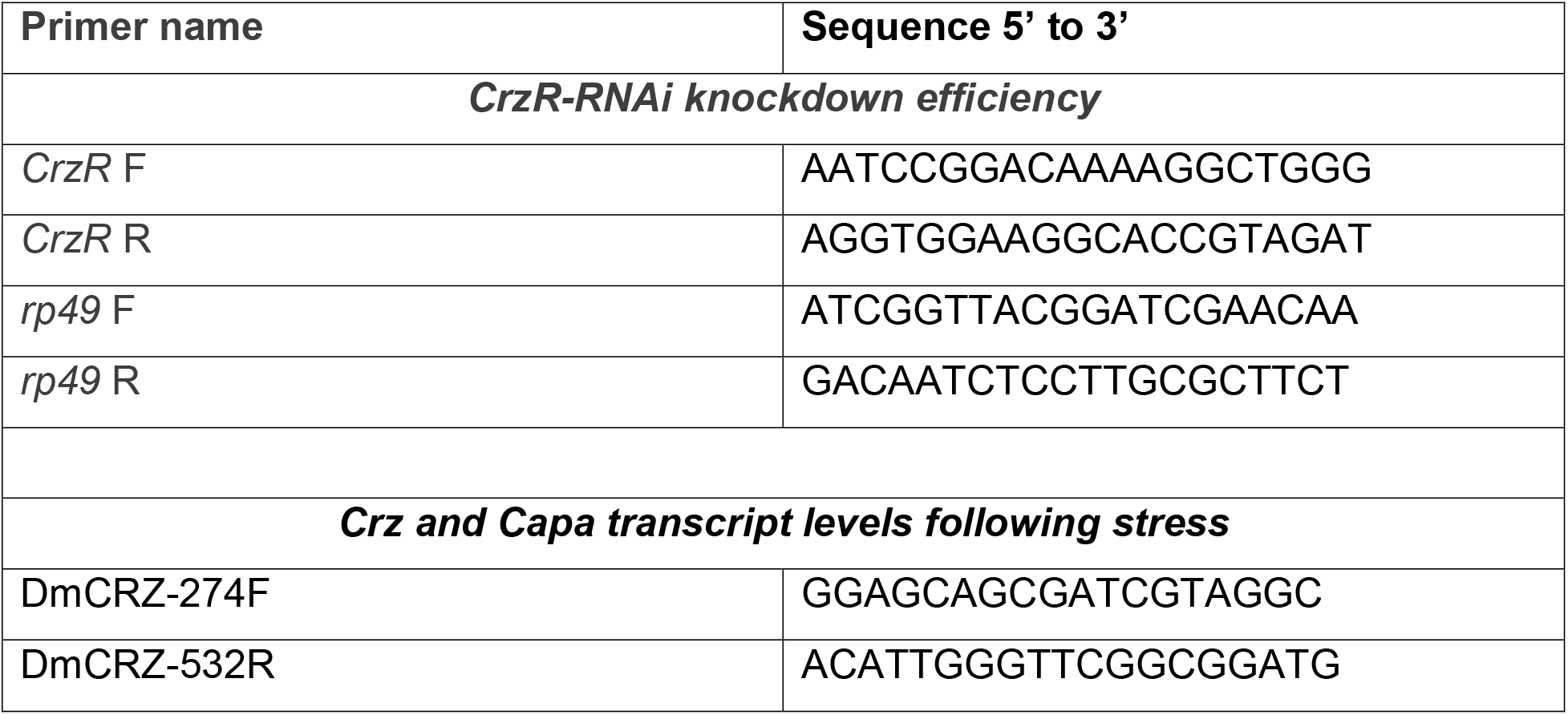
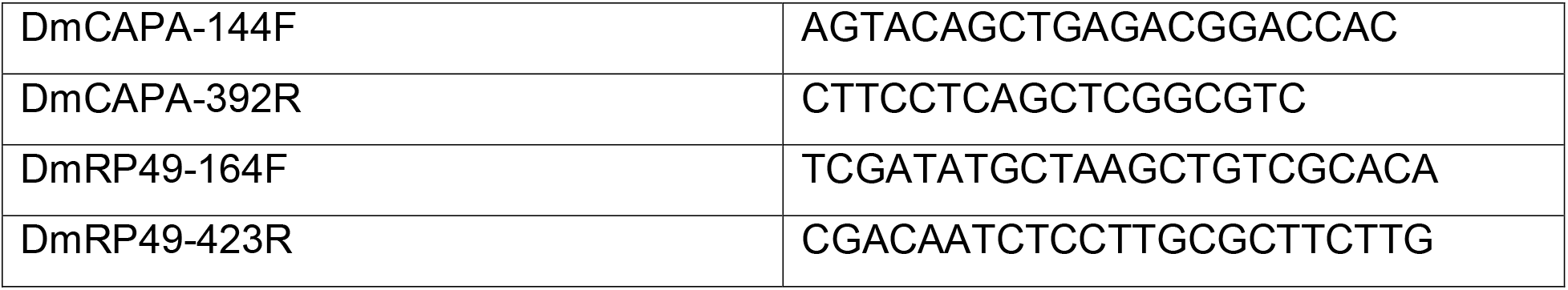
Primers used for qPCR.

Expression profiles were determined using three technical replicates per target and a minimum of three biological replicates. Finally, no reverse transcriptase control and no template control reactions were also conducted to verify primer fidelity and exclude false positives.

### Capillary feeding (CAFE) assay

A modified capillary feeding (CAFE) assay was used to monitor food intake of individual flies (10, 94). Capillaries were loaded with food comprised of 5% sucrose, 2% yeast extract and 0.1% propionic acid. Food consumption was measured daily and the cumulative food intake over 4 days was calculated. The experiment consisted of at least three biological replicates and 10 flies per replicate for each genotype.

### Defecation assay

Five to seven days old male flies were placed for one day on standard medium containing 0.05% bromophenol blue. Afterwards, prior to the evening activity phase, the flies were cooled down on ice, and then individually transferred to small glass tubes (diameter 5 mm) which were closed on one end with 2% agarose containing 4% sucrose and 0.05% bromophenol blue and a rubber plug to minimize dessication. The other end was closed by a small sponge. The tubes with flies were then placed horizontally in a special holder at 20°C and constant darkness in a climate chamber. Light was provided from below via an evenly illuminated red LED plate (λmax = 635 nm). Continuous optogenetic activation was achieved by a blue LED light source (λmax = 470 nm, about 0.6 mW/cm2) with a pulse of 60 seconds on/300 seconds off. Flies were recorded for 22h via a sCMOS camera placed directly above (DMK33UX178 and 22BUC03, ImagingSource, Bremen, Germany) at a frequency of 1 frame/15 min. Next, the lookup-table was inverted and the first frame was subtracted from all other frames using the <difference> operation in Fiji (80). Then, the number of feces droplets/22h was manually counted.

### Water content

To quantify water content following TRPA1-based thermogenetic activation, six day old male flies were kept on normal food for 24h at 18°C. All crosses were reared at 18°C even before isolation of flies for experiments. Approximately one hour before the experiments, groups of 15 flies (in six experimental replicates) were transferred to a 15mL conical tube to complete the ‘treatment’ incubation, whereby the tube containing the group of flies was transferred to 29°C (or kept at 18°C for control treatment) for a duration of 60 min. After the treatment duration at either 29°C or 18°C, flies were immediately frozen in liquid nitrogen and then thawed at room temperature for 1 hour and their wet weight was determined using a Sartorius Quintix124-1S analytical balance (Sartorius Canada Inc., Oakville, ON). The flies were then dried for 48 hrs at 60°C before their dry weight was recorded. Water content of flies was determined by subtracting the dry weight of the group of flies from the wet weights measured following the experiment.

### Mining public datasets for expression of genes

FlyAtlas2 database was mined to determine the distribution of *CrzR* in various tissues (95). The expression of *CAPAR* in the different regions of the gut and its cell types was determined using Flygut-*seq* (https://flygut.epfl.ch/) (96). A single-cell transcriptome atlas of the adult *Drosophila* brain (37) and VNC (36) was mined using SCope (http://scope.aertslab.org) to examine coexpression of *Crz* and *Capa*.

### Statistical analyses

#### Statistical Analyses for Peptide Injection Experiments

The peptide injection experiments (chill-coma recovery and desiccation survival) were each repeated on two-three different occasions (all within one month for each experiment). To account for this, time point was included as a factor in the models and was found to be significant in all cases. This suggests that the overall level of response differed among the different time points. The results as presented, show the treatment effects after controlling for the time effect. These analyses were performed in R (v. 3.4.1) (97) using two-way ANOVA and a Cox proportional hazards model (package survival), respectively.

#### Other Experiments

The experimental data presented in bar graphs represent means ± s.e.m. For the data presented in box-and-whisker plots, each individual value has been plotted and the horizontal line represents the median. Unless stated otherwise, one-way analysis of variance (ANOVA) followed by Tukey’s multiple comparisons test was used for comparisons between three genotypes and an unpaired *t*-test was used for comparisons between two genotypes. Stress survival data were compared using log-rank test, Mantel-Cox. All statistical analyses were performed using GraphPad Prism and the confidence intervals are included in the figure captions. The imaging data were analysed by a Shapiro-Wilk normality test followed by a paired Wilcoxon signed rank test in R.

## Acknowledgements

The authors would like to thank Dr. Olga Kubrak and Dr. Shreyas Jois for initial technical help and advice with experimental design. We would also like to thank Dr. Gilad Barnea (Brown University) for facilitating some of the experiments performed during the revision of this manuscript. We are grateful to the Bloomington Drosophila Stock Center (NIH P40OD018537), the Vienna Drosophila Resource Center, and Drs. Ian Orchard, Reinhard Predel, Jan Veenstra, Mark Brown, Heinrich Dircksen, Cornelis Grimmelikhuijzen, Stefan Thor, Jae Park and Shireen Davies for providing flies and reagents. Stina Höglund and the Imaging Facility at Stockholm University (IFSU) are acknowledged for maintenance of the confocal microscopes.

## Supplementary figures

**S1 Figure: Crz expression in the CNS of adult *Drosophila*.** *Crz*^*2*^-*GAL4* driven GFP and Crz-immunoreactivity is present in **(A)** dorsal lateral peptidergic neurons (DLPs) in the brain and **(B)** two to **(B^1^)** three pairs of male-specific neurons in the abdominal ganglia. **(C)** *Crz^1^>20X GFP* expression in the brain. Note that the Crz neuron arborizations can be seen in the antennal lobe (position marked with AL).

**S2 Figure: *Crz RNAi* knockdown efficiency and GeneSwitch manipulations (A-B)** *Crz^1^-GAL4* driven *Crz-RNAi (1)* but not *Crz-RNAi (2)* causes a significant decrease in anti-Crz staining (corrected total cell fluorescence, CTCF) in the brains of adult *Drosophila* (**** p < 0.0001 as assessed by One-way ANOVA). **(C)** Adult-specific pan-neuronal knockdown of *Crz* results in increased survival under ionic stress. “+” indicates RU486 fed flies and “-“ indicates control flies. Data are presented as survival curves (**** p < 0.0001, as assessed by Log-rank (Mantel-Cox) test).

**S3 Figure: *CrzR* is expressed in efferent neurons innervating the rectum. (A)** *CrzR-GAL4* driven GFP is not expressed in the midgut, hindgut and Malpighian tubules (MTs). **(B)** *CrzR* is weakly expressed in neurons innervating the rectum. White arrows indicate GFP staining in the region, presumably the water-resorbing rectal pads, which does not stain for phalloidin. **(C)** In adults, *CrzR* is not expressed in tissues (hindgut, MTs and rectal pad) associated with ionic and osmotic homeostasis (Data assembled from FlyAtlas 2)

**S4 Figure: Crz neuron processes superimpose those of *CrzR* expressing cells in the adult *Drosophila* CNS. (A, B)** Crz-producing dorsal lateral peptidergic neurons (DLPs) send projections to the pars intercerebralis near a set of *CrzR>GFP* expressing median neurosecretory cells. **(C)** Crz interneuron projections partly overlap *CrzR>GFP* expressing processes in the lateral horn. In **A-C**, note high *CrzR* expression (based on GFP intensity) in the local interneurons innervating the antennal lobe. **(D)** *CrzR-GAL4* drives strong GFP expression in three pairs of neurons in abdominal ganglion. Weak GFP expression is also observed in two pairs of neurons (indicated by a white arrow).

**S5 Figure: *CrzR* is expressed in DH44 neurons. (A)** *CrzR-GAL4* drives GFP expression in the median neurosecretory cells (MNCs) expressing diuretic hormone 44 (DH44) neuropeptide. **(B)** Higher magnification image of the DH44 MNCs (white box in **A**). Note that the GFP expression in DH44 neurons is weak and variable.

**S6 Figure: *CrzR* is not expressed in DH31 and Lk neurons.** *CrzR-GAL4* driven GFP is not expressed in diuretic hormone 31 (DH31) neurons of **(A, B)** the brain and **(C)** the ventral nerve cord. **B** shows a higher magnification image of the DH31 neurons surrounding the antennal lobe MNCs (white box in **A).** *CrzR-GAL4* driven GFP is not expressed in leucokinin (Lk) neurons of **(D)** the lateral horn (LHLKs), **(E)** subesophageal zone (SELKs) and **(F)** the abdominal ganglia (ABLKs).

**S7 Figure: *CrzR* expression in other peptidergic neurons.** *CrzR* is not expressed in **(A)** brain neurosecretory cells expressing *Drosophila* insulin-like peptide 2 (DILP2) and **(B)** adipokinetic hormone (AKH)-producing cells of the corpora cardiaca (CC). *CrzR-GAL4* drives GFP expression in **(C)** Hugin neurons and **(D)** neurons expressing FMRFamide-related peptide in the SEZ. Note that the exact identity of the neuropeptide present in the neurons labelled with FMRFamide antibody is unknown as it cross-reacts with multiple neuropeptides **(E, F)** *CrzR-GAL4* drives GFP expression in sLNv clock neurons labeled with anti-pigment dispersing factor (PDF) antibody. *CrzR>GFP* expression is present in the characteristic dorsal-projecting (DT) axons of the sLNvs.

**S8 Figure: Anatomical interaction between Crz and CAPA neurons. (A)** Crz interneurons in the abdominal ganglion send axon projections in close proximity to Va neurons. *CrzR-GAL4* drives GFP expression in CAPA/pyrokinin (CAPA/PK) producing neurons (labeled with anti-CAPA antibody) in **(B)** the subesophageal zone (SEZ) and **(C)** ventral nerve cord (VNC) of adult females. The three pairs of neurons in the VNC are referred to as Va neurons. **(D)** *Crz-GAL4* driven *trans-*Tango generates presynaptic signal (labeled with anti-GFP antibody) in the DLPs and a postsynaptic signal (labeled with anti-HA antibody) in various regions of the brain. **(E)** *Crz > trans-*Tango postsynaptic signal is absent in Va neurons of both males and females.

**S9 Figure: Peptide and transcript levels following stress and knockdown. (A)** Adult flies were either kept under normal conditions, starved, desiccated or rewatered (desiccated and then incubated on 1% aqueous agar) and Crz peptide levels monitored using immunohistochemistry. Crz peptide levels in all Crz neurons (see Figure 4A for representative images) are lower in starved, desiccated and rewatered flies compared to flies raised under normal conditions (* p < 0.05, ** p < 0.01 as assessed by One-way ANOVA). Starvation and refeeding do not impact **(B)** *Crz* and **(C)** *CAPA* transcript levels.

**S10 Figure: The *CAPA receptor (CAPAR)* is expressed in the adult gut and Malpighian tubules.** *CAPAR-GAL4* drives 20X GFP (*pJFRC81-10xUAS-Syn21-myr::GFP-p10*) expression in the adult **(A)** principal cells of the Malpighian tubules as well as in gut muscles. Note the lack of GFP staining in star-shaped stellate cells. **(B)** A schematic of the adult gut and heat map showing expression of *CAPAR* in different regions of the gut (R1 to R5) and its various cell types (VM, visceral muscle; EEC, enteroendocrine cell; EC, enterocyte; EB, enteroblast; ISC, intestinal stem cell; Ep, epithelium. Data was mined using Flygut-*seq*. The *CAPAR-GAL4* expression pattern is in agreement with the transcriptomic data.

**S11 Figure: *w*^*1118*^ > *TRPA1* control for water content and *CrzR* knockdown verification. (A)** There is no difference in the water content of *w*^*1118*^ > *TRPA1* flies incubated at 18°C and 29°C. **(B)** *Actin-GAL4* driven *CrzR-RNAi* (two independent RNAi lines) results in efficient knockdown of *CrzR* transcript in whole adult flies compared to control flies (*Actin>w*^*1118*^) as tested by qPCR. (** p < 0.01 as assessed by One-way ANOVA). **(C)** *Actin-GAL4* driven *CrzR-RNAi* has no impact on *Capa* transcript levels compared to control flies (*Actin>w*^*1118*^).

**S12 Figure: *Va-GAL4* and *CAPA-GAL4* drive GFP expression in CAPA neurons. (A)** *Va-GAL4* drives GFP expression in several neurons in the central nervous system, including a pair of CAPA/pyrokinin producing neurons in the SEZ and **(B)** Va neurons in the VNC (labeled with anti-CAPA antibody). **(C)** *CAPA-GAL4* does not drive GFP expression in the CAPA/pyrokinin neurons in the SEZ but it does so in Va neurons in the VNC **(D)**. Only 5 neurons are visible in this preparation; however, there are usually 6 neurons in most preparations **(D^1^)**. Note that the posterior-most pair of Va neurons send axonal projections into the abdominal nerve (indicated by the white arrow) where they terminate to form neurohemal release sites. The anterior two pairs send axons to a neurohemal plexus in the dorsal neural sheath.

**S13 Figure: *CrzR-GAL4* driven *CrzR-RNAi* influences osmotic and ionic stresses.** *CrzR-GAL4* driven *CrzR-RNAi(1)* results in **(A)** increased survival under desiccation and **(B)** ionic stress but has no impact on **(C)** chill-coma recovery. Data are presented as survival curves (**** p < 0.0001, as assessed by Log-rank (Mantel-Cox) test).

**S14 Figure: *Va-GAL4* driven *CrzR-RNAi* impacts food intake.** *CrzR* knockdown in Va neurons results in reduced cumulative food intake measured with CAFE assay over 4 days (*** p < 0.001, **** p < 0.0001 as assessed by One-way ANOVA).

## References

1. Owusu-Ansah E, Perrimon N. Stress signaling between organs in metazoa. Annu Rev Cell Dev Biol. 2015;31:497–522.

2. Droujinine IA, Perrimon N. Defining the interorgan communication network: systemic coordination of organismal cellular processes under homeostasis and localized stress. Front Cell Infect Microbiol. 2013;3:82.

3. Robertson GL, Shelton RL, Athar S. The osmoregulation of vasopressin. Kidney Int. 1976;10(1):25–37.

4. Verney E. TheAntidiuretic Hormone and the Factors Which Determine Its Release. Proc Roy Soc, London, s B. 1948;135:25–106.

5. Tatar M, Post S, Yu K. Nutrient control of *Drosophila* longevity. Trends Endocrinol Metab. 2014;25(10):509–17.

6. Nässel DR, Zandawala M. Recent advances in neuropeptide signaling in *Drosophila*, from genes to physiology and behavior. Prog Neurobiol. 2019;179:101607.

7. Rajan A, Perrimon N. *Drosophila* as a model for interorgan communication: lessons from studies on energy homeostasis. Dev Cell. 2011;21(1):29–31.

8. Fontana L, Partridge L, Longo VD. Extending healthy life span--from yeast to humans. Science. 2010;328(5976):321–6.

9. Kapan N, Lushchak OV, Luo J, Nässel DR. Identified peptidergic neurons in the *Drosophila* brain regulate insulin-producing cells, stress responses and metabolism by coexpressed short neuropeptide F and corazonin. Cell Mol Life Sci. 2012;69:4051–66.

10. Liu Y, Luo J, Carlsson MA, Nässel DR. Serotonin and insulin-like peptides modulate leucokinin-producing neurons that affect feeding and water homeostasis in *Drosophila*. J Comp Neurol. 2015;523(12):1840–63.

11. Nässel DR, Kubrak OI, Liu Y, Luo J, Lushchak OV. Factors that regulate insulin producing cells and their output in *Drosophila*. Front Physiol. 2013;4:252.

12. Nässel DR, Vanden Broeck J. Insulin/IGF signaling in Drosophila and other insects: factors that regulate production, release and post-release action of the insulin-like peptides. Cell Mol Life Sci. 2016;73(2): 271–90

13. Isabel G, Martin JR, Chidami S, Veenstra JA, Rosay P. AKH-producing neuroendocrine cell ablation decreases trehalose and induces behavioral changes in *Drosophila*. Am J Physiol Regul Integr Comp Physiol. 2005;288(2):R531–8.

14. Kubrak OI, Lushchak OV, Zandawala M, Nässel DR. Systemic corazonin signalling modulates stress responses and metabolism in *Drosophila*. Open Biol. 2016;6(11).

15. Lee G, Park JH. Hemolymph sugar homeostasis and starvation-induced hyperactivity affected by genetic manipulations of the adipokinetic hormone-encoding gene in *Drosophila melanogaster*. Genetics. 2004;167(1):311–23.

16. Zhao Y, Bretz CA, Hawksworth SA, Hirsh J, Johnson EC. Corazonin neurons function in sexually dimorphic circuitry that shape behavioral responses to stress in *Drosophila*. PLoS One. 2010;5(2):e9141.

17. Oh Y, Lai JS, Mills HJ, Erdjument-Bromage H, Giammarinaro B, Saadipour K, et al. A glucose-sensing neuron pair regulates insulin and glucagon in *Drosophila*. Nature. 2019;574(7779):559–64.

18. Ahmad M, He L, Perrimon N. Regulation of insulin and adipokinetic hormone/glucagon production in flies. Wiley Interdiscip Rev Dev Biol. 2020;9(2):e360.

19. Jourjine N, Mullaney BC, Mann K, Scott K. Coupled Sensing of Hunger and Thirst Signals Balances Sugar and Water Consumption. Cell. 2016;166(4):855–66.

20. Galikova M, Dircksen H, Nässel DR. The thirsty fly: Ion transport peptide (ITP) is a novel endocrine regulator of water homeostasis in *Drosophila*. PLoS Genet. 2018;14(8):e1007618.

21. Tian S, Zandawala M, Beets I, Baytemur E, Slade SE, Scrivens JH, et al. Urbilaterian origin of paralogous GnRH and corazonin neuropeptide signalling pathways. Sci Rep. 2016;6:28788.

22. Zandawala M, Tian S, Elphick MR. The evolution and nomenclature of GnRH-type and corazonin-type neuropeptide signaling systems. Gen Comp Endocrinol. 2018;264:64–77.

23. Veenstra JA. Does corazonin signal nutritional stress in insects? Insect Biochem Mol Biol. 2009;39(11):755–62.

24. Boerjan B, Verleyen P, Huybrechts J, Schoofs L, De Loof A. In search for a common denominator for the diverse functions of arthropod corazonin: a role in the physiology of stress? Gen Comp Endocrinol. 2010;166(2):222–33.

25. Miyamoto T, Slone J, Song X, Amrein H. A fructose receptor functions as a nutrient sensor in the *Drosophila* brain. Cell. 2012;151(5):1113–25.

26. Terhzaz S, Teets NM, Cabrero P, Henderson L, Ritchie MG, Nachman RJ, et al. Insect capa neuropeptides impact desiccation and cold tolerance. Proc Natl Acad Sci U S A. 2015;112(9):2882–7.

27. MacMillan HA, Nazal B, Wali S, Yerushalmi GY, Misyura L, Donini A, et al. Anti-diuretic activity of a CAPA neuropeptide can compromise *Drosophila* chill tolerance. J Exp Biol. 2018;221(Pt 19).

28. Lee G, Kim KM, Kikuno K, Wang Z, Choi YJ, Park JH. Developmental regulation and functions of the expression of the neuropeptide corazonin in *Drosophila melanogaster*. Cell and tissue research. 2008;331(3):659–73.

29. Tayler TD, Pacheco DA, Hergarden AC, Murthy M, Anderson DJ. A neuropeptide circuit that coordinates sperm transfer and copulation duration in *Drosophila*. Proc Natl Acad Sci U S A. 2012;109(50):20697–702.

30. Osterwalder T, Yoon KS, White BH, Keshishian H. A conditional tissue-specific transgene expression system using inducible GAL4. Proc Natl Acad Sci U S A. 2001;98(22):12596–601.

31. Kean L, Cazenave W, Costes L, Broderick KE, Graham S, Pollock VP, et al. Two nitridergic peptides are encoded by the gene capability in *Drosophila melanogaster*. Am J Physiol Regul Integr Comp Physiol. 2002;282(5):R1297–307.

32. Santos JG, Pollak E, Rexer KH, Molnar L, Wegener C. Morphology and metamorphosis of the peptidergic Va neurons and the median nerve system of the fruit fly, *Drosophila melanogaster*. Cell and tissue research. 2006;326(1):187–99.

33. Wegener C, Reinl T, Jansch L, Predel R. Direct mass spectrometric peptide profiling and fragmentation of larval peptide hormone release sites in *Drosophila melanogaster* reveals tagma-specific peptide expression and differential processing. J Neurochem. 2006;96(5):1362–74.

34. Zandawala M, Marley R, Davies SA, Nässel DR. Characterization of a set of abdominal neuroendocrine cells that regulate stress physiology using colocalized diuretic peptides in Drosophila. Cell Mol Life Sci. 2018;75(6):1099–115.

35. Imura E, Shimada-Niwa Y, Nishimura T, Huckesfeld S, Schlegel P, Ohhara Y, et al. The Corazonin-PTTH Neuronal Axis Controls Systemic Body Growth by Regulating Basal Ecdysteroid Biosynthesis in *Drosophila melanogaster*. Curr Biol. 2020.

36. Allen AM, Neville MC, Birtles S, Croset V, Treiber CD, Waddell S, et al. A single-cell transcriptomic atlas of the adult *Drosophila* ventral nerve cord. Elife. 2020;9.

37. Davie K, Janssens J, Koldere D, De Waegeneer M, Pech U, Kreft L, et al. A Single-Cell Transcriptome Atlas of the Aging *Drosophila* Brain. Cell. 2018;174(4):982–98 e20.

38. Bader R, Colomb J, Pankratz B, Schrock A, Stocker RF, Pankratz MJ. Genetic dissection of neural circuit anatomy underlying feeding behavior in *Drosophila*: distinct classes of hugin-expressing neurons. J Comp Neurol. 2007;502(5):848–56.

39. Helfrich-Forster C. The period clock gene is expressed in central nervous system neurons which also produce a neuropeptide that reveals the projections of circadian pacemaker cells within the brain of *Drosophila melanogaster*. Proc Natl Acad Sci U S A. 1995;92(2):612–6.

40. Schoofs A, Hückesfeld S, Schlegel P, Miroschnikow A, Peters M, Zeymer M, et al. Selection of motor programs for suppressing food intake and inducing locomotion in the *Drosophila* brain. PLoS biology. 2014;12(6):e1001893.

41. Schlegel P, Texada MJ, Miroschnikow A, Schoofs A, Huckesfeld S, Peters M, et al. Synaptic transmission parallels neuromodulation in a central food-intake circuit. Elife. 2016;5.

42. Melcher C, Pankratz MJ. Candidate gustatory interneurons modulating feeding behavior in the *Drosophila* brain. PLoS biology. 2005;3(9):e305.

43. Renn SC, Park JH, Rosbash M, Hall JC, Taghert PH. A pdf neuropeptide gene mutation and ablation of PDF neurons each cause severe abnormalities of behavioral circadian rhythms in *Drosophila*. Cell. 1999;99(7):791–802.

44. Nitabach MN, Taghert PH. Organization of the *Drosophila* circadian control circuit. Curr Biol. 2008;18(2):R84–93.

45. Sgourakis NG, Bagos PG, Papasaikas PK, Hamodrakas SJ. A method for the prediction of GPCRs coupling specificity to G-proteins using refined profile Hidden Markov Models. BMC Bioinformatics. 2005;6:104.

46. Choi YJ, Lee G, Hall JC, Park JH. Comparative analysis of Corazonin-encoding genes (Crz’s) in *Drosophila* species and functional insights into Crz-expressing neurons. J Comp Neurol. 2005;482(4):372–85.

47. Talay M, Richman EB, Snell NJ, Hartmann GG, Fisher JD, Sorkac A, et al. Transsynaptic Mapping of Second-Order Taste Neurons in Flies by trans-Tango. Neuron. 2017;96(4):783–95 e4.

48. Davies SA, Huesmann GR, Maddrell SH, O’Donnell MJ, Skaer NJ, Dow JA, et al. CAP2b, a cardioacceleratory peptide, is present in *Drosophila* and stimulates tubule fluid secretion via cGMP. Am J Physiol. 1995;269(6 Pt 2):R1321–6.

49. Halberg KA, Terhzaz S, Cabrero P, Davies SA, Dow JA. Tracing the evolutionary origins of insect renal function. Nat Commun. 2015;6:6800.

50. Yeoh JGC, Pandit AA, Zandawala M, Nassel DR, Davies SA, Dow JAT. DINeR: Database for Insect Neuropeptide Research. Insect Biochem Mol Biol. 2017;86:9–19.

51. Rodan AR, Baum M, Huang CL. The *Drosophila* NKCC Ncc69 is required for normal renal tubule function. Am J Physiol Cell Physiol. 2012;303(8):C883–94.

52. Paluzzi JP, Russell WK, Nachman RJ, Orchard I. Isolation, cloning, and expression mapping of a gene encoding an antidiuretic hormone and other CAPA-related peptides in the disease vector, Rhodnius prolixus. Endocrinology. 2008;149(9):4638–46.

53. Quinlan MC, Tublitz NJ, O’Donnell MJ. Anti-diuresis in the blood-feeding insect *Rhodnius prolixus* Stal: the peptide CAP2b and cyclic GMP inhibit Malpighian tubule fluid secretion. J Exp Biol. 1997;200(Pt 17):2363–7.

54. Terhzaz S, Cabrero P, Robben JH, Radford JC, Hudson BD, Milligan G, et al. Mechanism and function of *Drosophila* capa GPCR: a desiccation stress-responsive receptor with functional homology to human neuromedinU receptor. PLoS One. 2012;7(1):e29897.

55. Allan DW, St Pierre SE, Miguel-Aliaga I, Thor S. Specification of neuropeptide cell identity by the integration of retrograde BMP signaling and a combinatorial transcription factor code. Cell. 2003;113(1):73–86.

56. Davies SA, Cabrero P, Povsic M, Johnston NR, Terhzaz S, Dow JA. Signaling by *Drosophila* capa neuropeptides. Gen Comp Endocrinol. 2013;188:60–6.

57. Paluzzi JP. Anti-diuretic factors in insects: the role of CAPA peptides. Gen Comp Endocrinol. 2012;176(3):300–8.

58. Pollock VP, McGettigan J, Cabrero P, Maudlin IM, Dow JA, Davies SA. Conservation of capa peptide-induced nitric oxide signalling in Diptera. J Exp Biol. 2004;207(Pt 23):4135–45.

59. Sajadi F, Curcuruto C, Al Dhaheri A, Paluzzi JV. Anti-diuretic action of a CAPA neuropeptide against a subset of diuretic hormones in the disease vector *Aedes aegypti*. J Exp Biol. 2018;221(Pt 7).

60. Sajadi F, Uyuklu A, Paputsis C, Lajevardi A, Wahedi A, Ber LT, et al. CAPA neuropeptides and their receptor form an anti-diuretic hormone signaling system in the human disease vector, *Aedes aegypti*. Sci Rep. 2020;10(1):1755.

61. Koyama T, Terhzaz S, Naseem MT, Nagy S, Rewitz K, Dow JAT, et al. A nutrient-responsive hormonal circuit controls energy and water homeostasis in *Drosophila*. bioRxiv. 2020:2020.07.24.219592.

62. Schooley DA, Horodyski FM, Coast GM. 9 - Hormones Controlling Homeostasis in Insects. In: Gilbert LI, editor. Insect Endocrinology. San Diego: Academic Press; 2012. p. 366–429.

63. Bergland AO, Chae HS, Kim YJ, Tatar M. Fine-scale mapping of natural variation in fly fecundity identifies neuronal domain of expression and function of an aquaporin. PLoS Genet. 2012;8(4):e1002631.

64. Miyamoto T, Amrein H. Diverse roles for the *Drosophila* fructose sensor Gr43a. Fly (Austin). 2014;8(1):19–25.

65. Fujii S, Yavuz A, Slone J, Jagge C, Song X, Amrein H. Drosophila sugar receptors in sweet taste perception, olfaction, and internal nutrient sensing. Curr Biol. 2015;25(5):621–7.

66. Root CM, Ko KI, Jafari A, Wang JW. Presynaptic facilitation by neuropeptide signaling mediates odor-driven food search. Cell. 2011;145(1):133–44.

67. Ko KI, Root CM, Lindsay SA, Zaninovich OA, Shepherd AK, Wasserman SA, et al. Starvation promotes concerted modulation of appetitive olfactory behavior via parallel neuromodulatory circuits. eLife. 2015;4.

68. Martelli C, Pech U, Kobbenbring S, Pauls D, Bahl B, Sommer MV, et al. SIFamide Translates Hunger Signals into Appetitive and Feeding Behavior in Drosophila. Cell Rep. 2017;20(2):464–78.

69. Yu Y, Huang R, Ye J, Zhang V, Wu C, Cheng G, et al. Regulation of starvation-induced hyperactivity by insulin and glucagon signaling in adult *Drosophila*. Elife. 2016;5.

70. Bharucha KN, Tarr P, Zipursky SL. A glucagon-like endocrine pathway in *Drosophila* modulates both lipid and carbohydrate homeostasis. J Exp Biol. 2008;211(Pt 19):3103–10.

71. Inagaki HK, Panse KM, Anderson DJ. Independent, reciprocal neuromodulatory control of sweet and bitter taste sensitivity during starvation in *Drosophila*. Neuron. 2014;84(4):806–20.

72. Tian S, Egertova M, Elphick MR. Functional Characterization of Paralogous Gonadotropin-Releasing Hormone-Type and Corazonin-Type Neuropeptides in an Echinoderm. Front Endocrinol (Lausanne). 2017;8:259.

73. Sha K, Choi SH, Im J, Lee GG, Loeffler F, Park JH. Regulation of ethanol-related behavior and ethanol metabolism by the Corazonin neurons and Corazonin receptor in *Drosophila melanogaster*. PLoS One. 2014;9(1):e87062.

74. Asahina K, Watanabe K, Duistermars BJ, Hoopfer E, Gonzalez CR, Eyjolfsdottir EA, et al. Tachykinin-expressing neurons control male-specific aggressive arousal in *Drosophila*. Cell. 2014;156(1-2):221–35.

75. Choi YJ, Lee G, Park JH. Programmed cell death mechanisms of identifiable peptidergic neurons in *Drosophila melanogaster*. Development. 2006;133(11):2223–32.

76. Nicholson L, Singh GK, Osterwalder T, Roman GW, Davis RL, Keshishian H. Spatial and temporal control of gene expression in *Drosophila* using the inducible GeneSwitch GAL4 system. I. Screen for larval nervous system drivers. Genetics. 2008;178(1):215–34.

77. Pfeiffer BD, Truman JW, Rubin GM. Using translational enhancers to increase transgene expression in *Drosophila*. Proc Natl Acad Sci U S A. 2012;109(17):6626–31.

78. Chen TW, Wardill TJ, Sun Y, Pulver SR, Renninger SL, Baohan A, et al. Ultrasensitive fluorescent proteins for imaging neuronal activity. Nature. 2013;499(7458):295–300.

79. Shafer OT, Kim DJ, Dunbar-Yaffe R, Nikolaev VO, Lohse MJ, Taghert PH. Widespread receptivity to neuropeptide PDF throughout the neuronal circadian clock network of *Drosophila* revealed by real-time cyclic AMP imaging. Neuron. 2008;58(2):223–37.

80. Schindelin J, Arganda-Carreras I, Frise E, Kaynig V, Longair M, Pietzsch T, et al. Fiji: an open-source platform for biological-image analysis. Nat Methods. 2012;9(7):676–82.

81. Pollak E, Eckert M, Molnar L, Predel R. Differential sorting and packaging of capa-gene related products in an insect. J Comp Neurol. 2005;481(1):84–95.

82. Veenstra JA, Davis NT. Localization of corazonin in the nervous system of the cockroach *Periplaneta americana*. Cell and tissue research. 1993;274(1):57–64.

83. Veenstra JA, Agricola HJ, Sellami A. Regulatory peptides in fruit fly midgut. Cell and tissue research. 2008;334(3):499–516.

84. Kaufmann C, Brown MR. Adipokinetic hormones in the African malaria mosquito, *Anopheles gambiae*: Identification and expression of genes for two peptides and a putative receptor. Insect Biochem Molec. 2006;36(6):466–81.

85. Grimmelikhuijzen CJP. FMRFamide is generally occuring in the nervous system of coelenterates. Histochemistry. 1983;78:361–81.

86. Dircksen H, Zahnow CA, Gaus G, Keller R, Rao KR, Riehm JP. The ultrastructure of nerve endings containing pigmentdispersing hormone (PDH) in crustacean sinus glands: identification by an antiserum against synthetic PDH. Cell and tissue research. 1987;250:377–87.

87. Nässel DR, Cantera R, Karlsson A. Neurons in the cockroach nervous system reacting with antisera to the neuropeptide leucokinin I. J Comp Neurol. 1992;322(1):45–67.

88. Cabrero P, Radford JC, Broderick KE, Costes L, Veenstra JA, Spana EP, et al. The Dh gene of *Drosophila melanogaster* encodes a diuretic peptide that acts through cyclic AMP. J Exp Biol. 2002;205(Pt 24):3799–807.

89. Park D, Veenstra JA, Park JH, Taghert PH. Mapping peptidergic cells in *Drosophila*: where DIMM fits in. PLoS One. 2008;3(3):e1896.

90. Wahedi A, Gade G, Paluzzi JP. Insight Into Mosquito GnRH-Related Neuropeptide Receptor Specificity Revealed Through Analysis of Naturally Occurring and Synthetic Analogs of This Neuropeptide Family. Front Endocrinol (Lausanne). 2019;10:742.

91. Folk DG, Han C, Bradley TJ. Water acquisition and partitioning in Drosophila melanogaster: effects of selection for desiccation-resistance. J Exp Biol. 2001;204(Pt 19):3323–31.

92. Feng Y, Ueda A, Wu CF. A modified minimal hemolymph-like solution, HL3.1, for physiological recordings at the neuromuscular junctions of normal and mutant *Drosophila* larvae. J Neurogenet. 2004;18(2):377–402.

93. Livak KJ, Schmittgen TD. Analysis of relative gene expression data using real-time quantitative PCR and the 2(-Delta Delta C(T)) Method. Methods. 2001;25(4):402–8.

94. Ja WW, Carvalho GB, Mak EM, de la Rosa NN, Fang AY, Liong JC, et al. Prandiology of *Drosophila* and the CAFE assay. Proc Natl Acad Sci U S A. 2007;104(20):8253–6.

95. Leader DP, Krause SA, Pandit A, Davies SA, Dow JAT. FlyAtlas 2: a new version of the *Drosophila melanogaster* expression atlas with RNA-Seq, miRNA-Seq and sex-specific data. Nucleic Acids Res. 2018;46(D1):D809–D15.

96. Dutta D, Dobson AJ, Houtz PL, Glasser C, Revah J, Korzelius J, et al. Regional Cell-Specific Transcriptome Mapping Reveals Regulatory Complexity in the Adult Drosophila Midgut. Cell Rep. 2015;12(2):346–58.

97. R Core Team. R: A language and environment for statistical computing. . R Foundation for Statistical Computing, Vienna, Austria 2017;URL https://www.R-project.org/.

